# Local Protein Dynamics during Microvesicle Exocytosis in Neuroendocrine Cells

**DOI:** 10.1101/245647

**Authors:** Agila Somasundaram, Justin Taraska

## Abstract

Calcium triggered exocytosis is key to many physiological processes, including neurotransmitter and hormone release by neurons and endocrine cells. Dozens of proteins regulate exocytosis, yet the temporal and spatial dynamics of these factors during vesicle fusion remain unclear. Here we use total internal reflection fluorescence microscopy to visualize local protein dynamics at single sites of exocytosis of small synaptic-like microvesicles in live cultured neuroendocrine PC12 cells. We employ two-color imaging to simultaneously observe membrane fusion (using vesicular acetylcholine transporter (VAChT) tagged to pHluorin) and the dynamics of associated proteins at the moments surrounding exocytosis. Our experiments show that many proteins, including the SNAREs syntaxin1 and VAMP2, the SNARE modulator tomosyn, and Rab proteins, are pre-clustered at fusion sites and rapidly lost at fusion. The ATPase NSF is locally recruited at fusion. Interestingly, the endocytic BAR domain-containing proteins amphiphysin1, syndapin2, and endophilins are dynamically recruited to fusion sites, and slow the loss of vesicle membrane-bound cargo from fusion sites. A similar effect on vesicle membrane protein dynamics was seen with the over-expression of the GTPases dynamin1 and dynamin2. These results suggest that proteins involved in classical clathrin-mediated endocytosis can regulate exocytosis of synaptic-like microvesicles. Our findings provide insights into the dynamics, assembly, and mechanistic roles of many key factors of exocytosis and endocytosis at single sites of microvesicle fusion in live cells.

## INTRODUCTION

Exocytosis is the cellular process in which cytoplasmic membrane-bound vesicles fuse with the plasma membrane and release their contents into the extracellular space. During synaptic transmission, action potentials depolarize the presynaptic terminal triggering Ca^2+^ influx into the cell. Local elevations in intracellular Ca^2+^ cause synaptic vesicles (SVs) in the terminal to fuse with the plasma membrane, releasing neurotransmitters into the synaptic cleft (Jahn and Fasshauer, 2012). SV exocytosis is a carefully orchestrated process that involves multiple steps and dozens of proteins. SVs are small, ∼ 50 nm in diameter, and contain a repertoire of proteins on their membranes (Takamori *et al*., 2006). These vesicular membrane proteins and several cytoplasmic and plasma membrane-associated proteins play important roles in regulating SV exocytosis (Sudhof, 2013d).

Specifically, SNARE proteins are thought to drive SV fusion with the plasma membrane (Sudhof and Rothman, 2009). The vesicular SNARE, synaptobrevin (VAMP), and the plasma membrane SNAREs, syntaxin and SNAP-25, form a four-helical zippered complex that pulls the two lipid bilayers together resulting in fusion. The SNAREs are sufficient for fusion in vitro (van den Bogaart *et al*., 2010). However, under physiological conditions, several proteins such as Rabs and their effector molecules (Fukuda, 2008), complexin (Trimbuch and Rosenmund, 2016), the Ca sensor synaptotagmin (Sudhof, 2013a), tomosyn (Ashery *et al*., 2009; Bielopolski *et al*., 2014), CAPS (Stevens and Rettig, 2009), Munc18 and Munc13 (Sudhof and Rothman, 2009) have been proposed to regulate the steps leading to fusion, including docking (attaching the SV to the active zone) and priming (preparing the SV for fusion) (Sudhof, 2013d). While much is known about the biochemical properties of these proteins and their physiological effects, a comprehensive understanding of their spatial and temporal dynamics during SV exocytosis in live cells is lacking, partly due to the small size and the challenges associated with labeling and imaging single vesicles in synaptic terminals (Kavalali and Jorgensen, 2014). Understanding the dynamics of these key mediators of SV exocytosis will provide direct insights into their regulatory functions, biological mechanisms, and roles in disease.

Neuroendocrine PC12 cells have two distinct calcium triggered exocytic vesicle pools, ∼50 nm diameter synaptic-like microvesicles (SLMVs) and the larger ∼150 nm diameter dense core vesicles (DCVs) (Thomas-Reetz and De Camilli, 1994). Both vesicle populations fuse with the plasma membrane with a rise in intracellular calcium. However, each vesicle type exhibits distinct calcium sensitivities and fusion kinetics, and is responsible for the release of different signals (Ninomiya *et al*., 1997). Specifically, DCVs release peptide neurotransmitters, proteins, and amines while microvesicles release the chemical neurotransmitter acetylcholine or γ-aminobutyric acid. Furthermore, SLMVs are stimulated at much lower intracellular calcium concentrations allowing for a more rapid burst of fusion during depolarization (Ninomiya *et al*., 1997). SLMVs are structurally and functionally similar to neuronal SVs (Thomas-Reetz and De Camilli, 1994). They maintain an acidic pH, accumulate neurotransmitters, and fuse with the plasma membrane to release their luminal contents in a Ca^2+^-dependent manner. Furthermore, they contain many of the same proteins required for cargo packaging, transport, exocytosis, and recycling, but unlike SVs, SLMVs don’t appear clustered at active zones via synapsins (Thomas-Reetz and De Camilli, 1994). Aside from the interest in their signaling functions in endocrine cells, these vesicles have been used as experimentally tractable surrogates for the study of SV behavior (de Wit *et al*., 2001; Brauchi *et al*., 2008; Sochacki *et al*., 2012). Here, we used total internal reflection fluorescence (TIRF) microscopy to monitor the dynamics of over two dozen proteins during calcium-triggered exocytosis of single SLMVs in PC12 cells (Sochacki *et al*., 2012; Trexler *et al*., 2016). In an imaging-based screen examining key exocytic and endocytic proteins, we show that many proteins, including syntaxin, VAMP, tomosyn, and Rab GTPases are pre-clustered at fusion sites and rapidly diffuse away following exocytosis. Interestingly, BAR domain-containing proteins, and dynamin, known to be important in clathrin-mediated endocytosis, are recruited during fusion and influence the loss of vesicle membrane proteins from the fusion sites. Our study provides insights into the local dynamics, assembly, and function of key regulators of exocytosis and endocytosis during microvesicle fusion in live cells.

## RESULTS

To image microvesicle fusion in PC12 cells, we expressed the vesicular acetylcholine transporter (VAChT) tagged on its luminal side to pHluorin, a pH-sensitive variant of green fluorescent protein (Miesenbock *et al*., 1998). VAChT is targeted specifically to synaptic-like microvesicles (SLMVs) in PC12 cells (Liu and Edwards, 1997), and VAChT-pHluorin has been used to track SLMV exocytosis (Brauchi *et al*., 2008; Sochacki *et al*., 2012). pHluorin fluorescence is quenched in the acidic lumen of the vesicle, but upon fusion the lumenal pH is neutralized by the extracellular buffer causing pHluorin signal to dramatically increase, enabling the detection of exocytic events. To stimulate exocytosis, we depolarized cells by applying buffer containing high extracellular KCl using a superfusion pipette positioned close to the cell (Trexler *et al*., 2016; Trexler and Taraska, 2017). Membrane depolarization caused a substantial increase in the frequency of fusion events as shown in Supplemental Figure 1A. Fusion events were detected as sudden bright flashes of green fluorescence (Figure 1A). This is observed as a local sharp increase in signal that decays with time as VAChT-pH diffuses away from the sites of exocytosis (Figure 1A, bottom, and Supplemental Figure 1B). In the red fluorescent channel, we monitored co-expressed proteins fused to mCherry, mRFP, or mKate2 (Table 1). In the example shown in Figure 1, VAChT-pH was co-expressed with mRFP-Rab3A (Figure 1B), where measurable changes in signal were detected during fusion (Figure 1B, bottom). Background-subtracted fluorescence intensities from hundreds of individual fusion events were extracted, normalized, and averaged in the green and red channels for every protein examined in the study to produce the average time-dependent changes in local protein signals at fusion sites (Figure 1, C and D). We analyzed ∼ 8,000 fusion events from over 300 cells to track, quantitate, and characterize the dynamics of dozens of proteins (Table 1). To verify that fusion events predominantly occurred from microvesicles in our experimental system, we stimulated PC12 cells co-expressing neuropeptide Y (NPY) tagged to GFP (to label large dense core vesicles (LDCVs) and VAChT-pHuji (Shen *et al*., 2014; Martineau *et al*., 2017). As expected, VAChT-labeled fusion events were detected in the pHuji channel (23 events in 4 cells), but not in the NPY-GFP channel, and no increase in GFP signal was measured in regions corresponding to VAChT-labeled events (not shown). These results further confirm that VAChT specifically targets SLMVs in PC12 cells.

**Figure 1.**
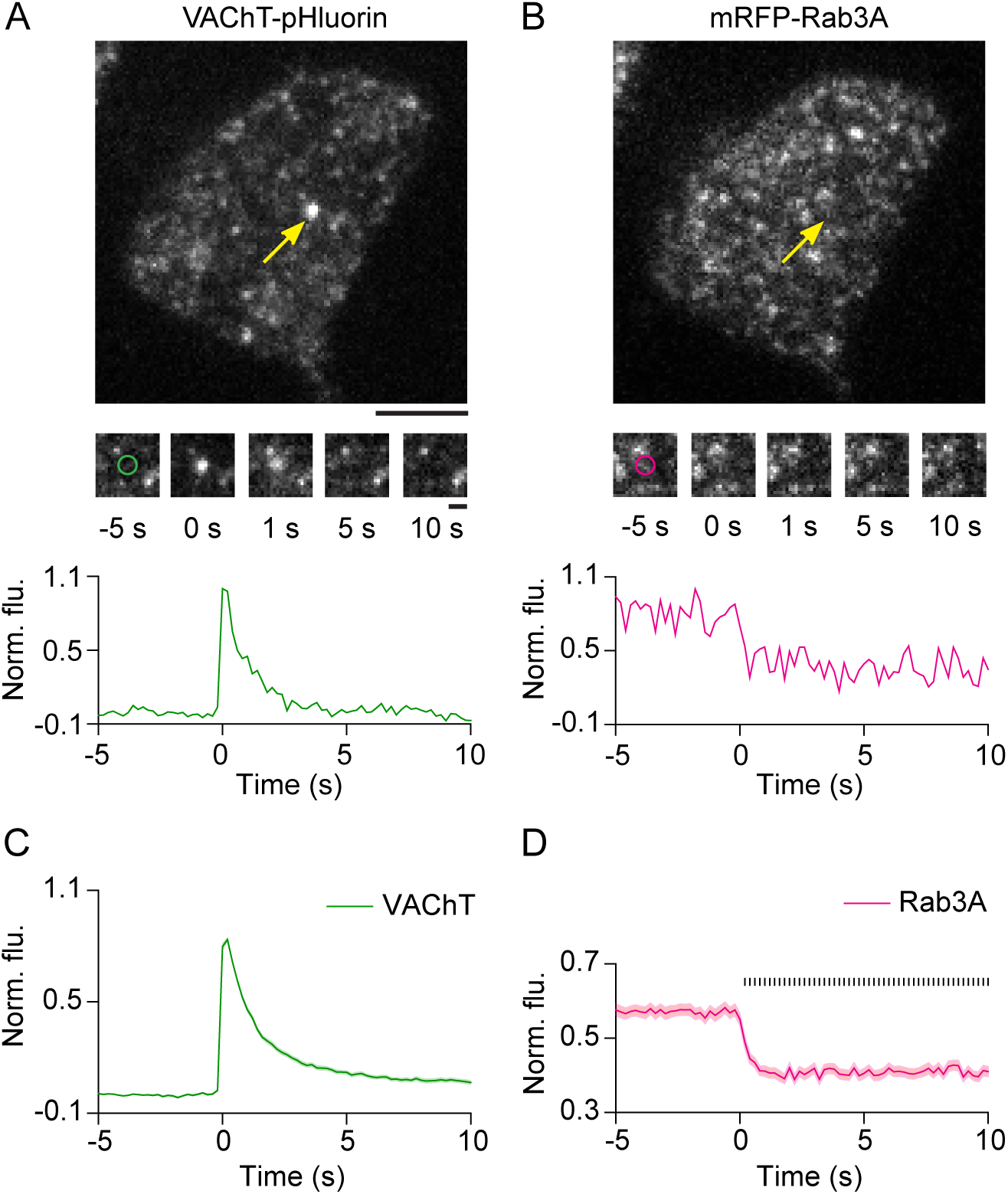
Imaging protein dynamics at SLMV fusion sites in PC12 cells. (A, B) Image of a PC12 cell transfected with VAChT-pH (A) and mRFP-Rab3A (B) imaged using TIRF microscopy. Arrows (yellow) show a fusion event in the green channel and the corresponding region in the red channel, after application of stimulation buffer. Bar, 5 μm. (Middle) Snapshots of the fusion event shown above at the indicated time-points. ‘0 s’ indicates the manually identified first frame of brightening in the green channel. Circles (∼ 1 µm diameter) represent regions used for intensity analysis. Bar, 1 μm. (Bottom) Time-lapse traces of normalized fluorescence intensities for the event shown above in the green and red channels. (C, D) Average time-lapse traces of normalized fluorescence intensities for VAChT-pH (green) and mRFP-Rab3A (magenta) (196 events, 5 cells). Individual event traces were time-aligned to 0 s, which corresponds to the fusion frame in the green channel. Small vertical black lines in (D) indicate p < 0.05 (paired Student’s t-test) when comparing every time-point after 3 s with the average pre-fusion value obtained from −5 to −3 s. Standard errors are plotted as shaded areas around the average traces.

### Rab GTPases and effectors are rapidly lost from sites of SLMV exocytosis

We first examined the dynamics of the Rab family of GTPases and Rab effector molecules, which play important roles in targeting and docking synaptic vesicles to the plasma membrane (Fukuda, 2008). In PC12 cells, Rab27A and Rab3A were localized at exocytic sites before fusion and diffused away rapidly following exocytosis, consistent with their vesicle membrane-anchored nature (Figure 2, A and B, and Supplemental Figure 2). The cytosolic Rab3A effector molecule, Rabphilin3A, also displayed similar localization and behavior (Figure 2C and Supplemental Figure 2). An early endosomal Rab, Rab5A (Woodman, 2000), showed some enrichment at fusion sites, that slowly decreased following fusion (Figure 2D, Supplemental Figure 2). Rab27B did not exhibit specific localization at SLMVs or a change in intensity following fusion (Figure 2E and Supplemental Figure 2). These results demonstrate that the Rab proteins, Rab27A and Rab3A, and the effector Rabphilin3A, are targeted to microvesicle sites before exocytosis and are dynamically lost into the cytosol or plasma membrane following fusion.

**Figure 2.**
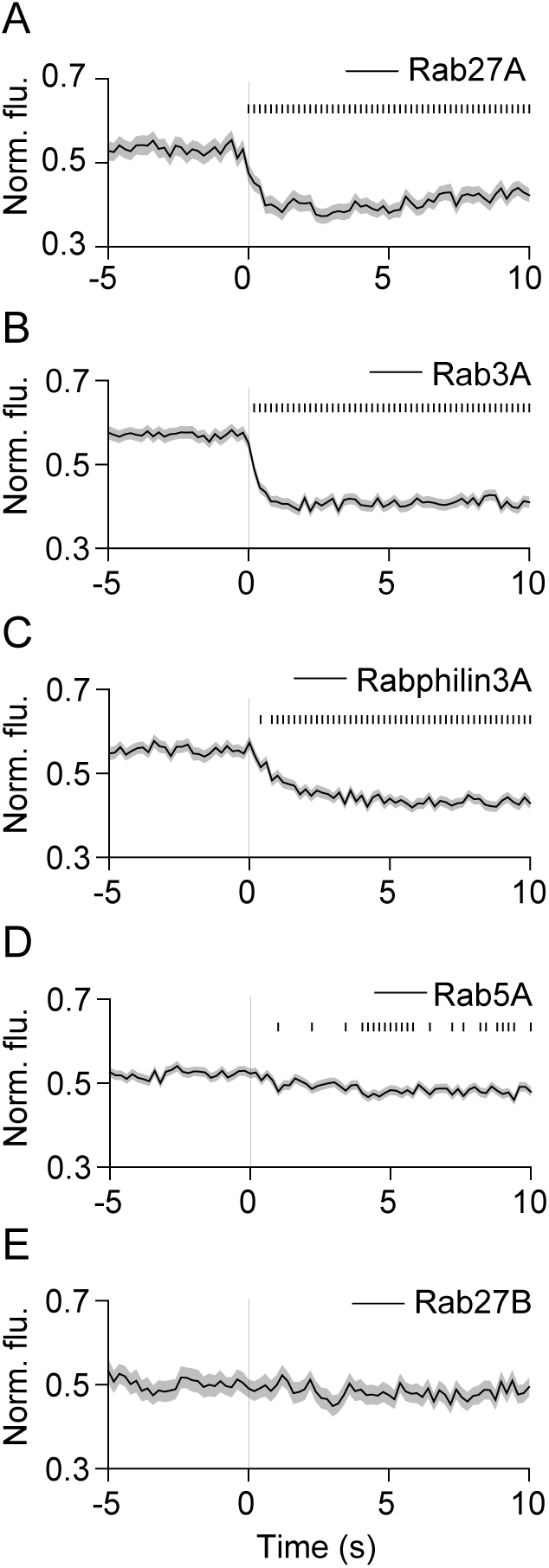
Dynamics of Rab proteins during SLMV fusion. (A-E) Average time-lapse traces of normalized fluorescence intensities for: (A) mCherry-Rab27A (103 events, 4 cells), (B) mRFP-Rab3A (196 events, 5 cells), (C) mCherry-Rabphilin3A (198 events, 6 cells), (D) mCherry**-** Rab27B (73 events, 3 cells), and (E) mCherry-Rab5A (276 events, 6 cells). Individual event traces were time-aligned to 0 s (vertical black line), which corresponds to the fusion frame in the green channel. Small vertical black lines indicate p < 0.05 (paired Student’s t-test) when comparing every time-point after 3 s with the average pre-fusion value obtained from −5 to −3 s. Standard errors are plotted as shaded areas around the average traces.

### SNAREs, syntaxin1 and VAMP2, are clustered at fusion sites and lost following fusion

Docked vesicles fuse with the plasma membrane by the concerted action of the SNARE proteins syntaxin, SNAP25 and VAMP2 (Sudhof and Rothman, 2009). We found that the plasma membrane SNARE, syntaxin1, decreased in intensity at fusion sites during exocytosis (Figure 3A). Similar to the distribution seen in LDCVs (Lang *et al*., 2001; Barg *et al*., 2010; Gandasi and Barg, 2014; Ullrich *et al*., 2015), radial scan analysis of syntaxin1 fluorescence revealed locally elevated syntaxin1 that diffused away following fusion, indicating that syntaxin1 is clustered on the plasma membrane at sites of microvesicle exocytosis (Supplemental Figure 3). We did not observe re-clustering of syntaxin1 at the original fusion sites (Supplemental Figure 3). The plasma membrane-attached SNARE, SNAP25, did not exhibit increased localization at fusion sites, or a substantial change in signal or distribution during fusion (Figure 3B and Supplemental Figure 3). VAMP2 showed some concentration at fusion sites, consistent with its expression on the vesicular membrane, and diffused away following fusion (Figure 3C and Supplemental Figure 3). To rule out potential artifacts induced by the red fluorescent tag, we examined the distribution and dynamics of cytosolic mCherry and membrane-attached farnesyl-mCherry during fusion. Cytosolic mCherry was diffusely distributed both before and during fusion, and did not show substantial changes in signal during fusion (Figure 3D and Supplemental Figure 3). Farnesyl-mCherry signal increased slightly during fusion, perhaps due the delivery of a small amount of probe contained on the vesicle to the plasma membrane (Figure 3E and Supplemental Figure 3). Thus, the control mCherry proteins exhibited small or no changes in signal during fusion, that are markedly different from the dynamics of syntaxin1 and VAMP2 described above. In conclusion, the SNAREs syntaxin1 and VAMP2 are locally pre-assembled at microvesicle fusion sites and diffuse away from the sites of release.

**Figure 3.**
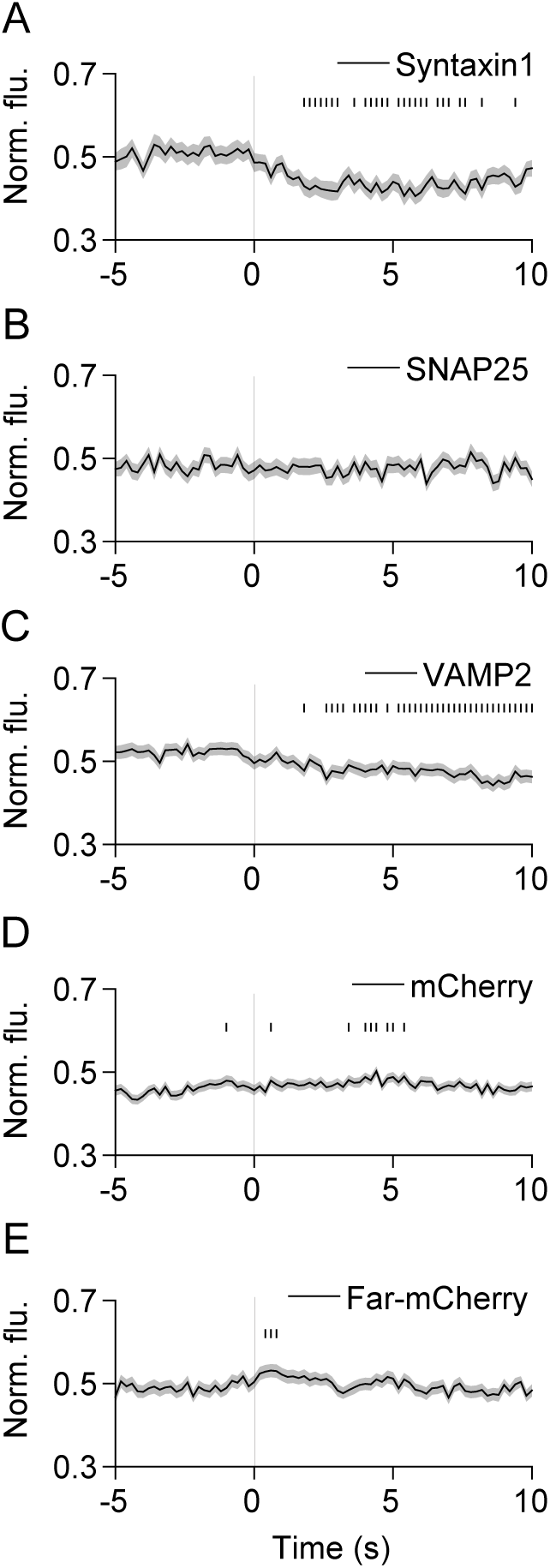
SNAREs syntaxin1 and VAMP2 diffuse away from sites of SLMV fusion. (A-E) Average time-lapse traces of normalized fluorescence intensities for: (A) dCMV-mCherry-syntaxin1 (76 events, 4 cells), (B) dCMV-mCherry-SNAP25 (97 events, 8 cells), (C) VAMP2-mCherry (172 events, 7 cells), (D) mCherry (274 events, 8 cells), and (E) farnesyl-mCherry (166 events, 9 cells). Individual event traces were time-aligned to 0 s (vertical black line), which corresponds to the fusion frame in the green channel. Small vertical black lines indicate p < 0.05 (paired Student’s t-test) when comparing every time-point after 3 s with the average pre-fusion value obtained from −5 to −3 s. Standard errors are plotted as shaded areas around the average traces.

### SNARE modulators exhibit diverse behaviors during SLMV fusion

SNARE-mediated fusion is regulated by several proteins (Sudhof, 2013d). To gain insights into the function of these SNARE modulators in live cells, we next examined their dynamics during SLMV fusion in PC12 cells. Multiple roles have been proposed for the small protein complexin, which is thought to either clamp the SNARE complex in a partially zippered state, or facilitate fusion (Brose, 2008; Yoon *et al*., 2008; An *et al*., 2010; Wragg *et al*., 2013). Complexin2 signal decreases slightly during fusion (Figure 4A); however, most of the protein appeared diffusely distributed before and after fusion (Supplemental Figure 5). A small decrease in signal was also seen with CAPS (Figure 4B), a protein essential for SV priming (Jockusch *et al*., 2007). Unlike complexin2, CAPS displayed some preferential localization at fusion sites at rest (Supplemental Figure 5). The Ca sensor synaptotagmin1 (Sudhof, 2013a) was concentrated at fusion sites but, surprisingly, did not diffuse away following fusion (Supplemental Figures 4A and 5). Tomosyn, thought to play a largely inhibitory role in SV priming (Gracheva *et al*., 2006; McEwen *et al*., 2006; Ashery *et al*., 2009; Bielopolski *et al*., 2014; Cazares *et al*., 2016) was located as clusters at vesicles and diffused away following fusion (Figure 4C and Supplemental Figure 5). Munc18a and Munc13, proteins proposed to bind the SNAREs and have essential roles in SV fusion (Sudhof and Rothman, 2009), did not exhibit significant changes in dynamics during fusion (Supplemental Figure 4, B and C, and Supplemental Figure 5). Interestingly, the ATPase NSF was transiently recruited to fusion sites during exocytosis (Figure 4D and Supplemental Figure 5), consistent with its role in disassembling SNARE complexes (Ryu *et al*., 2016). Overall, SNARE modulators exhibited diverse and subtle behaviors, with some pre-assembled at fusion sites and lost following fusion (CAPS and tomosyn), some recruited to fusion sites (NSF), and others localized at fusion sites throughout fusion (synaptotagmin1). These data likely reflect the diverse functions of these distinct classes of proteins along with the transient and step-wise nature of their activities during microvesicle fusion.

**Figure 4.**
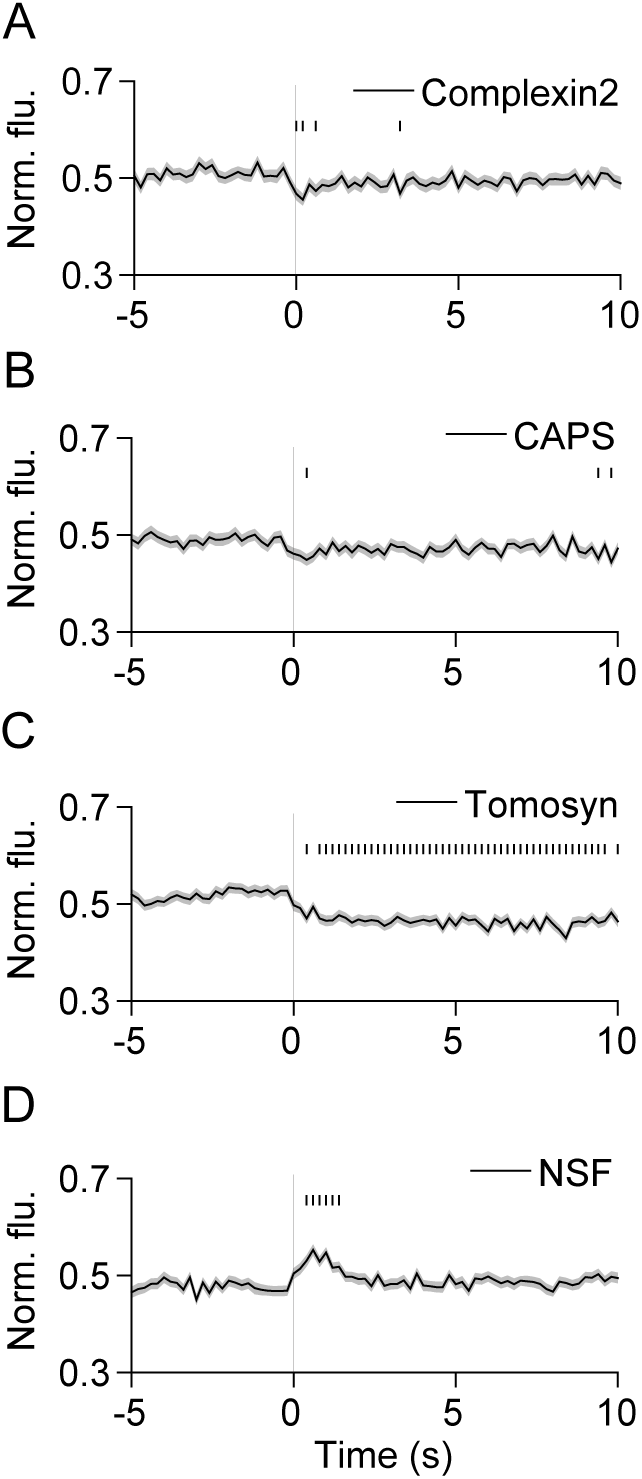
SNARE modulators exhibit diverse behaviors during SLMV fusion. (A-C) Average time-lapse traces of normalized fluorescence intensities for: (A) complexin2-mCherry (202 events, 7 cells), (B) CAPS-mKate2 (232 events, 5 cells), (C) mCherry-tomosyn (289 events, 9 cells), and (D) NSF-mCherry (274 events, 5 cells). Individual event traces were time-aligned to 0 s (vertical black line), which corresponds to the fusion frame in the green channel. Small vertical black lines indicate p < 0.05 (paired Student’s t-test) when comparing every time-point after 3 s with the average pre-fusion value obtained from −5 to −3 s. Standard errors are plotted as shaded areas around the average traces.

### BAR domain proteins are recruited to sites of SLMV exocytosis

Previous studies have shown that the proteins amphiphysin, syndapin, and dynamin, involved in the maturation and scission of clathrin-coated endocytic structures (McMahon and Boucrot, 2011; Daumke *et al*., 2014), are recruited to exocytic sites (Tsuboi *et al*., 2004; Jaiswal *et al*., 2009; Trexler *et al*., 2016), and regulate Ca^2+^-dependent cargo release from LDCVs in endocrine cells (Graham *et al*., 2002; Holroyd *et al*., 2002; Tsuboi *et al*., 2004; Min *et al*., 2007; Fulop *et al*., 2008; Llobet *et al*., 2008; Anantharam *et al*., 2010; Anantharam *et al*., 2011; Samasilp *et al*., 2012; Trexler *et al*., 2016), and constitutive secretion from post-Golgi vesicles (Jaiswal *et al*., 2009). Because SLMVs are about 5-fold smaller than LDCVs and likely have different curvatures, we hypothesized that a distinct set of proteins might be recruited (Xu and Xu, 2008; Zhang and Jackson, 2010). We first imaged the dynamics of the curvature sensing/inducing BAR domain proteins amphiphysin1 and syndapin2. Both proteins were specifically recruited to microvesicles during exocytosis (Figure 5 and Supplemental Figure 6A). BAR domain proteins endophilinA1 and endophilinA2 were also recruited during fusion (Supplemental Figure 7, A and B). EndophilinB1, however, appeared localized at fusion sites even at rest, and did not exhibit significant changes in dynamics during fusion (Supplemental Figure 7D). Overall, these results indicate that specific BAR domain proteins are recruited to SLMVs at exocytosis in neuroendocrine cells.

**Figure 5.**
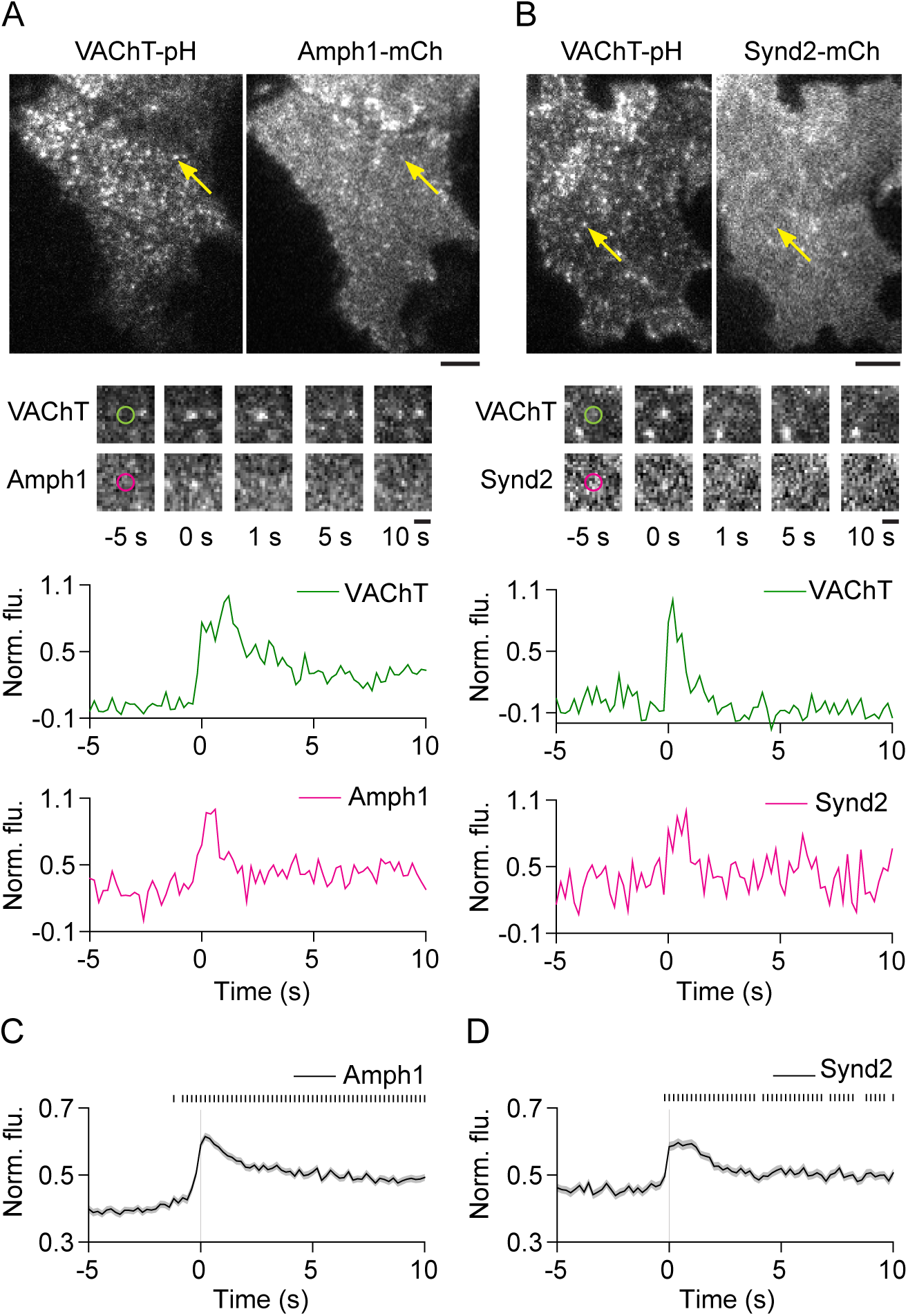
Amphiphysin1 and syndapin2 are transiently recruited to SLMV fusion sites. (A, B) Images of PC12 cells expressing (A) VAChT-pH and amphiphysin1-mCherry or (B) VAChT-pH and syndapin2-mCherry. Arrowheads show fusion events in the green channel and the corresponding regions in the red channel, after application of stimulation buffer. Bar, 5 μm. (Middle) Snapshots of the fusion event shown above, at the indicated time-points. Time-point ‘0 s’ indicates the first frame of brightening in the green channel. Bar, 1 μm. (Bottom) Time-lapse traces of normalized fluorescence intensities in the green and red channels for the fusion event shown above. (C, D) Average time-lapse traces of normalized fluorescence intensities in the red channel for (C) Amph1-WT (371 events, 9 cells) and (D) Synd2-WT (228 events, 6 cells). Individual event traces were time-aligned to 0 s (vertical black line), which corresponds to the fusion frame in the green channel. Small vertical black lines indicate p < 0.05 (paired Student’s t-test) when comparing every time-point after 3 s with the average pre-fusion value obtained from −5 to −3 s. Standard errors are plotted as shaded areas around the average traces.

To determine if amphiphysin1 and syndapin2 recruitment to the site of fusion is driven by the curvature-sensitive BAR domains (Daumke *et al*., 2014), we imaged mutants lacking BAR domains. Unlike the wild-type (WT) proteins, Amph1-ΔBAR and Synd2-ΔBAR did not show specific localization at fusion sites during exocytosis (Figure 6, A-D, and Supplemental Figure 6A), indicating that the BAR domain is required for amphiphysin1 and syndapin2 recruitment. Deletion of the protein-protein interaction domain (SH3) (David *et al*., 1996; Grabs *et al*., 1997; Qualmann *et al*., 1999) in amphiphysin1 (Amph1-ΔSH3) and syndapin2 (Synd2-ΔSH3) did not prevent recruitment (Figure 6, E and F, and Supplemental Figure 6A), indicating that the SH3 domain is not required for their localization to SLMVs during fusion. Moreover, the BAR domain of syndapin2 (Synd2-BAR) showed strong recruitment (Figure 6H and Supplemental Figure 6A), whereas over-expression of the SH3 domain alone of amphiphysin1 (Amph1-SH3) did not show recruitment to fusion sites (Figure 6G and Supplemental Figure 6A). Thus, amphiphysin1 and syndapin2 recruitment is dependent on BAR domains, suggesting that these proteins could be targeted to the highly curved neck of the expanding fusion pore. Interestingly, recruitment appeared to slightly precede fusion (Figure 6) suggesting a possible localization of these proteins to a transient hemi-fusion state (Zhao *et al*., 2016). Also, the retention of these proteins at fusion sites seconds after fusion (Figure 6 and Supplemental Figure 6A), suggests either the persistence of a curved neck of the fusion pore or the rapid formation of endocytic structures at or near the fusion sites (Watanabe *et al*., 2013a; Watanabe *et al*., 2013c). Radial scan analysis shows diffusional spread of VAChT-pH following fusion in the presence of Synd2-BAR (Supplemental Figure 6B), suggesting that at least some proteins can leave the sites of microvesicle exocytosis.

**Figure 6.**
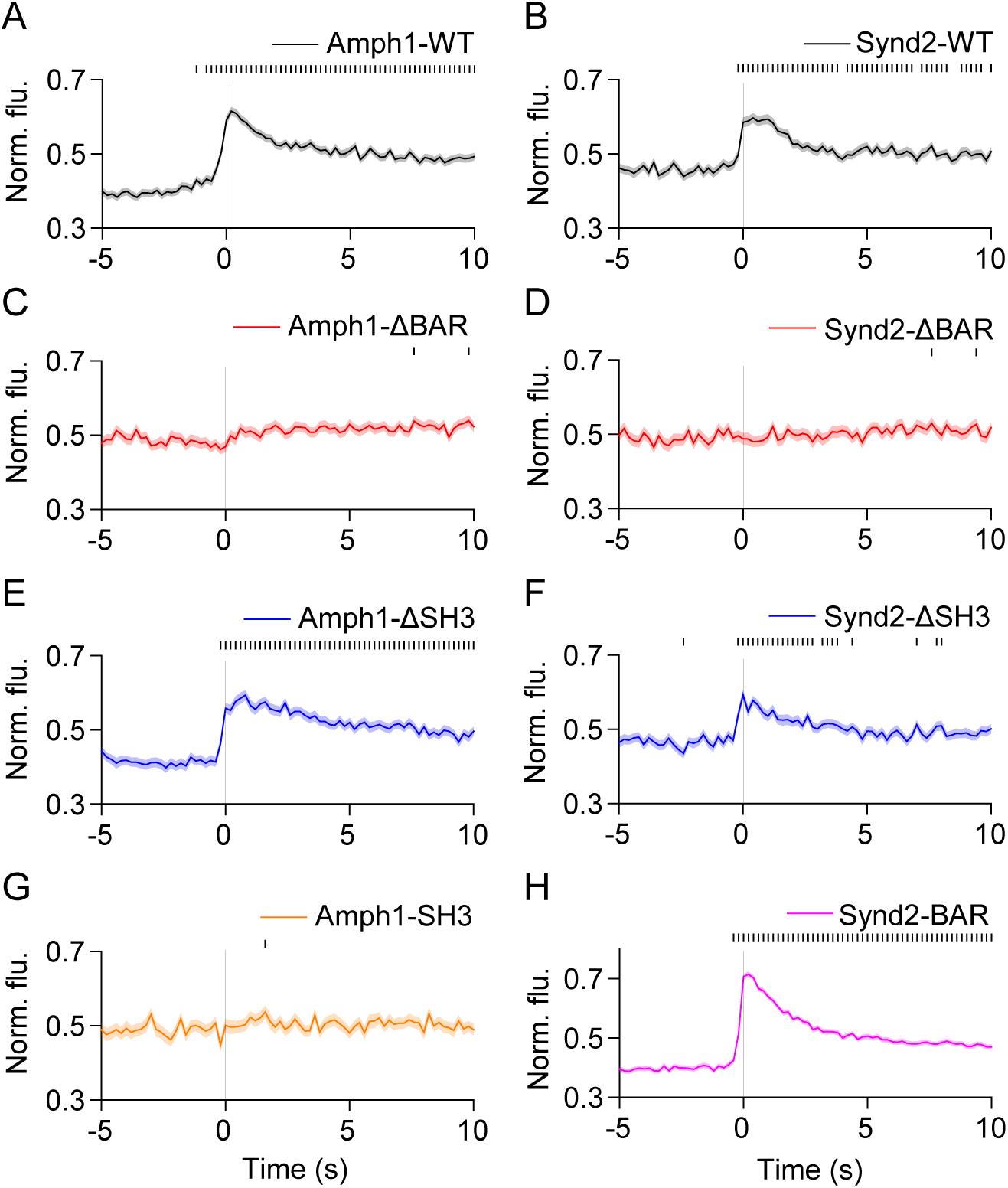
Amphiphysin1 and syndapin2 recruitment to SLMV fusion sites is dependent on the BAR domain. (A-H) Average time-lapse traces of normalized fluorescence intensities in the red channel for: (A) Amph1-WT (371 events, 9 cells), (B) Synd2-WT (228 events, 6 cells), (C) Amph1-ΔBAR (169 events, 4 cells), (D) Synd2-ΔBAR (136 events, 11 cells), (E) Amph1-ΔSH3 (214 events, 13 cells), (F) Synd2-ΔSH3 (184 events, 9 cells), (G) Amph1-SH3 (117 events, 8 cells), and (H) Synd2-BAR (418 events, 13 cells). Individual event traces were time-aligned to 0 s (vertical black line), which corresponds to the fusion frame in the green channel. Small vertical black lines indicate p < 0.05 (paired Student’s t-test) when comparing every time-point after 3 s with the average pre-fusion value obtained from −5 to −3 s. Standard errors are plotted as shaded areas around the average traces.

### BAR domain proteins modulate loss of SLMV membrane cargo from fusion sites

We next investigated whether the recruitment of BAR domain proteins described above impacts the loss of vesicle membrane cargo from fusion sites by measuring VAChT-pH fluorescence decay (Sochacki *et al*., 2012). Expression of Amph1-WT did not significantly alter VAChT-pH loss when compared with farnesyl-mCherry (Supplemental Figure 8A, τ = 2.39 ± 0.04 s vs. τ = 2.33 ± 0.04 s). However, over-expression of Amph1-ΔBAR and Amph1-SH3, proteins that were not recruited to fusion sites (Figure 6 and Supplemental Figure 6), resulted in faster VAChT-pH decay when compared with WT (Figure 7A, τ = 1.36 ± 0.06 s, τ = 1.88 ± 0.08 s). On the other hand, over-expression of Amph1-ΔSH3, which was recruited to fusion sites (Figure 6 and Supplemental Figure 6), resulted in slower VAChT-pH decay (Figure 7A, τ = 3.12 ± 0.05 s). These results suggest that amphiphysin1 recruitment during SLMV fusion slows the loss of vesicle membrane proteins from fusion sites.

**Figure 7.**
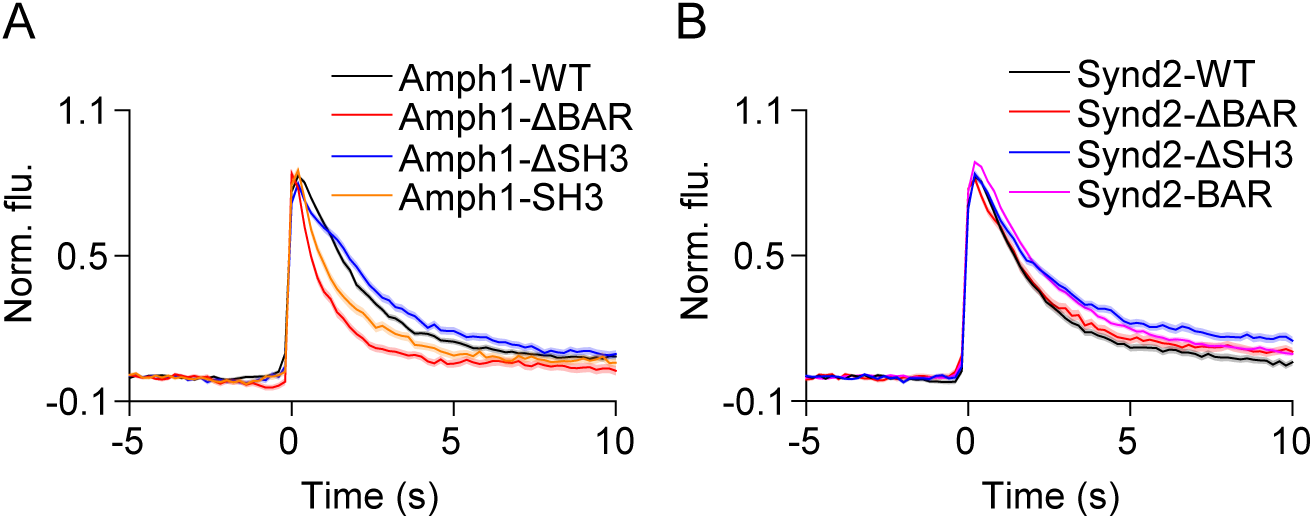
Amphiphysin1 and syndapin2 mutants slow the loss of VAChT-pH from fusion sites. (A) Average time-lapse traces of normalized VAChT-pH fluorescence intensities in PC12 cells co-expressing VAChT-pH and (A) WT or mutant amphiphysin1, and (B) WT or mutant syndapin2 constructs. Individual event traces were time-aligned to 0 s, which corresponds to the frame of fusion. Standard errors are plotted as shaded areas around the average traces.

We obtained similar results with syndapin2. Like Amph1-WT, expression of Synd2-WT did not alter VAChT-pH decay when compared with farnesyl-mCherry (Supplemental Figure 8B, τ = 2.28 ± 0.04 s vs. τ = 2.33 ± 0.04 s). Synd2-ΔBAR, which was not recruited to fusion sites (Figure 6 and Supplemental Figure 6), exhibited VAChT decay slightly slower than Synd2-WT (Figure 7B, τ = 2.63 ± 0.05). However, Synd2-ΔSH3 and Synd2-BAR, which showed strong and specific recruitment to fusion sites (Figure 6 and Supplemental Figure 6), substantially slowed down VAChT decay (Figure 7B, τ = 3.43 ± 0.07 s, τ = 2.94 ± 0.04 s). Thus, the lack of the SH3 domain in amphiphysin1 and syndapin2 further slows down the loss of membrane cargo when compared with WT (Figure 7). Furthermore, the BAR domain proteins endophilinA1, endophilinA2 and endophilinB1 that showed recruitment or localization at fusion sites (Supplemental Figure 7), resulted in significantly slower VAChT-pH decay when compared with farnesyl-mCherry (Supplemental Figure 8C, τ = 3.24 ± 0.04 s, τ = 3.07 ± 0.06 s, τ = 4.16 ± 0.07 s). Intriguingly, the VAChT-pH decay with farnesyl-mCherry was slower than with cytosolic mCherry (Supplemental Figure 8D, τ = 2.33 ± 0.04 s vs. 2.07 ± 0.10 s). It is possible that the small increase in farnesyl-mCherry seen during exocytosis (Figure 3 and Supplemental Figure 3) alters membrane properties including fluidity or packing to slow down cargo release (Stachowiak *et al*., 2012). Because we are interested in the impact of membrane-associated BAR domain proteins on fusion dynamics, we chose to compare the resulting VAChT-pH decay profiles with that seen with farnesyl-mCherry (Supplemental Figure 8). Overall, our results indicate that BAR domain proteins slow the loss of vesicle membrane cargo from fusion sites during SLMV exocytosis.

We further examined the effects of knock-down of endogenous amphiphysin1 and syndapin2 on VAChT-pH loss by treating cells with siRNA against these molecules. Western blot analysis revealed syndapin2 expression in PC12 cells, which was reduced with siRNA treatment (Supplemental Figure 9A). We were unsuccessful in our attempts to detect endogenous amphiphysin1 using commercial antibodies (data not shown). Contrary to our expectation, the VAChT-pH decay in PC12 cells treated with siSyndapin2 was slower than that with control siRNA (Supplemental Figure 9B, τ = 4.01 ± 0.06 s vs. τ = 3.15 ± 0.07 s). It is possible that syndapin2 knock-down resulted in increased recruitment of other BAR domain proteins to fusion sites, but this complication was not explored further. Nonetheless, our experiments with amphiphysin1 and syndapin2 mutants, and endophilins, provide evidence for the modulation of microvesicle membrane cargo dynamics by BAR domain proteins.

### Dynamin is recruited to fusion sites and delays loss of SLMV membrane cargo

Expression of amphiphysin1 and syndapin2 mutants lacking the SH3 domains resulted in slower VAChT-pH decay than that seen with full-length proteins (Figure 7), suggesting a role for SH3 binding partners in hastening the loss of VAChT from fusion sites. To test this, we examined the dynamics of the well-studied SH3 binding partner, the GTPase dynamin (David *et al*., 1996; Qualmann *et al*., 1999; Ferguson and De Camilli, 2012). During clathrin-mediated endocytosis (CME), dynamin localizes to the curved neck of the invaginating clathrin-coated pit through its interactions with BAR domain proteins and causes scission (Daumke *et al*., 2014). Dynamin1 and dynamin2 have also been shown to cluster at LDCV fusion sites (Holroyd *et al*., 2002; Tsuboi *et al*., 2004; Trexler *et al*., 2016), and regulate fusion pore expansion (Holroyd *et al*., 2002; Tsuboi *et al*., 2004; Min *et al*., 2007; Fulop *et al*., 2008; Anantharam *et al*., 2011; Samasilp *et al*., 2012; Gonzalez-Jamett *et al*., 2013; Fan *et al*., 2015; Trexler *et al*., 2016). We found that dynamin1 is transiently recruited to SLMVs during fusion (Supplemental Figure 10A), and slows down the loss of VAChT-pH when compared with control (Figure 8A, τ = 3.47 ± 0.05 s vs. τ = 2.33 ± 0.04 s). Dyn1-K44A, a GTPase mutant, was recruited to SLMV fusion sites (Supplemental Figure 10B), suggesting that the GTPase activity of dynamin1 is dispensable for its recruitment. Dyn1-K44A, however, hastened VAChT-pH decay (Figure 8B, τ = 2.09 ± 0.08 s). Thus, the recruitment of functional dynamin1 to SLMV fusion sites slows down loss of membrane cargo from the fusion site.

**Figure 8.**
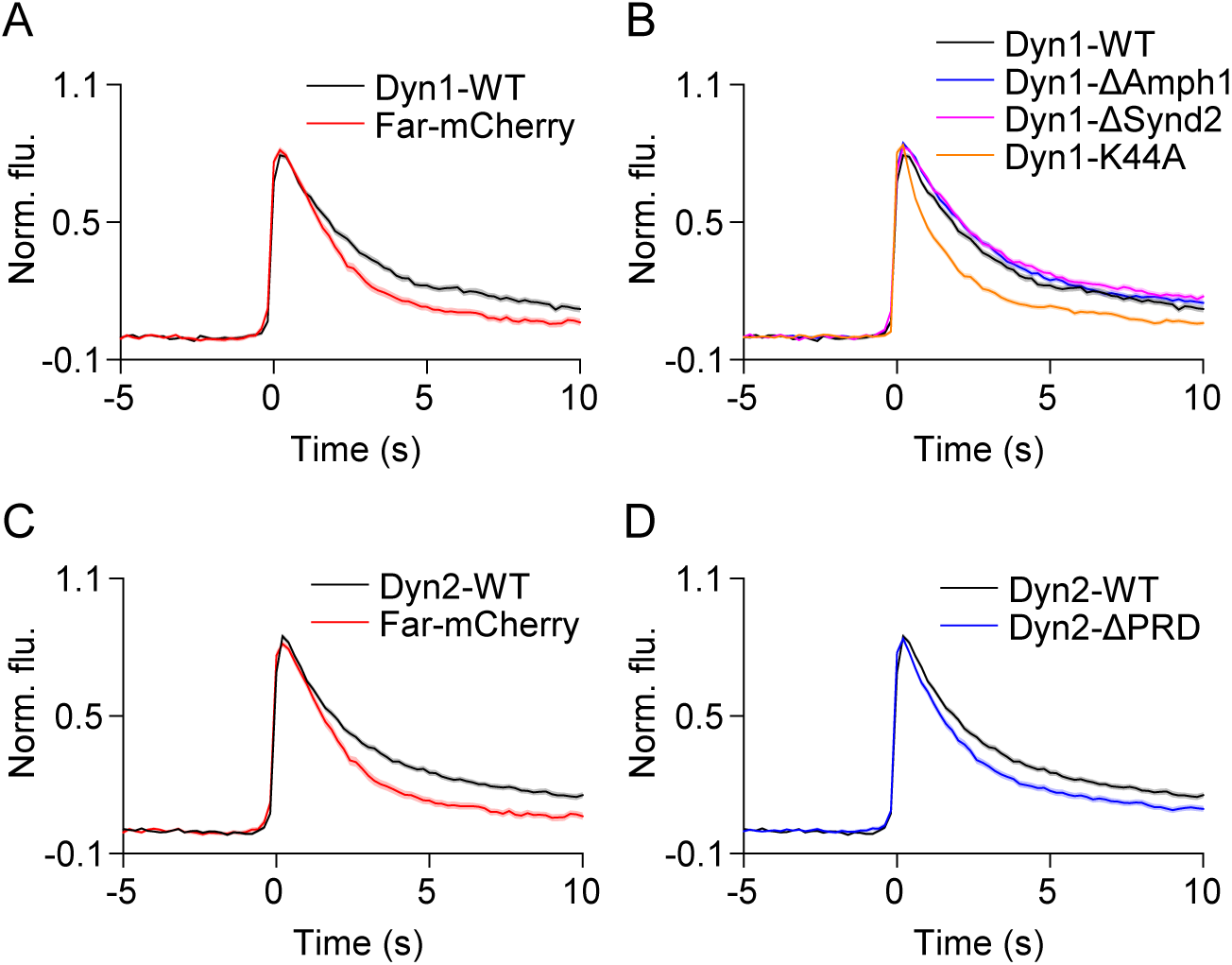
Dynamin1 and dynamin2 slow the loss of VAChT-pH from fusion sites. (A-D) Average time-lapse traces of normalized VAChT-pH fluorescence intensities in PC12 cells co-expressing VAChT-pH and (A-B) WT or mutant dynamin1, and (C-D) WT or mutant dynamin2 constructs. Individual event traces were time-aligned to 0 s, which corresponds to the frame of fusion. Standard errors are plotted as shaded areas around the average traces.

We next examined the effects of disrupting dynamin1’s interactions with BAR domain proteins by mutating residues in its proline rich domain (PRD) (Okamoto *et al*., 1997). Dyn1 deficient in binding amphiphysin1 (Dyn1-833-838A, Dyn1-ΔAmph1) (Grabs *et al*., 1997) did not show specific localization at fusion sites, whereas dynamin1 deficient in binding syndapin2 (Dyn1-S774E/S778E, Dyn1-ΔSynd2) (Anggono *et al*., 2006) was recruited to SLMVs during fusion (Supplemental Figure 10, C and D). Both mutants, however, resulted in VAChT-pH decay that was comparable to that observed with Dyn1-WT (Figure 8B, τ = 3.68 ± 0.05 s, τ = 3.93 ± 0.07 s), and slower than with farnesyl-mCherry. It is possible that background levels of expression of Dyn1-ΔAmph1 was sufficient to affect VAChT dynamics. Thus, our findings suggest that dynamin1 slows the loss of membrane cargo, and disrupting its interactions with either amphiphysin1 or syndapin2 does not lessen its effects.

Unlike dynamin1, dynamin2 (Dyn2-WT) did not show substantial recruitment to fusion sites (Supplemental Figure 10E). PC12 cells expressing Dyn2-WT, however, exhibited slower VAChT-pH decay when compared with farnesyl-mCherry (Figure 8C, τ = 3.56 ± 0.09 s vs. τ = 2.33 ± 0.04 s). This suggests that low levels of Dyn2-WT at fusion sites can affect cargo loss, or that dynamin2 acts at a step prior to or distant from exocytosis. Dynamin2 lacking the PRD domain (Dyn2-ΔPRD) (Supplemental Figure 10F) resulted in faster loss of membrane cargo (Figure 8D, τ = 2.69 ± 0.06 s), suggesting that dynamin2’s interaction with BAR domain proteins is essential for this effect. Taken together, these results suggest that dynamin1 and dynamin2 slow the loss of SLMV membrane cargo from fusion sites. This is consistent with the observation that Amph1-SH3, which sequesters SH3 binding partners such as dynamin (Shupliakov *et al*., 1997; Wigge *et al*., 1997; Holroyd *et al*., 2002), results in faster VAChT-pH decay (Figure 7A). These results also suggest that the slower VAChT-pH decay seen in Amph1-ΔSH3 and Synd2-ΔSH3 mutants (Figure 7) lacking dynamin binding is likely due to a lack of interactions with SH3 binding partners other than dynamin.

Examination of the dynamics of N-WASP, a protein that binds SH3-domain containing proteins and stimulates actin polymerization during CME (Qualmann *et al*., 1999; Kessels and Qualmann, 2002; Yamada *et al*., 2009), did not reveal significant changes in signal or distribution during fusion (Supplemental Figure 11A), or a substantial change in the VAChT-pH decay when compared with control (τ = 2.64 ± 0.05 s, vs. τ = 2.33 ± 0.04 s). Interestingly, we observed a slow, but significant increase in clathrin around fusion sites several seconds after fusion (Supplemental Figure 11B). Moreover, apart from the specific and transient recruitment of amphiphysin1 to fusion sites described earlier (Figure 5), in 5 out of 9 cells we observed a marked increase in amphiphysin1 clusters across the plasma membrane that peaked tens of seconds after stimulation and then disappeared (Supplemental Figure 12), suggesting compensatory endocytosis of released cargo or membrane, controlled by amphiphysin. This additional slow recruitment did not seem correlated to fusion sites. Lastly, the adaptor protein AP-2 and the scaffolding protein intersectin did not show significant recruitment to fusion sites (Supplemental Figure 11, C and D), suggesting perhaps the presence of sufficient amounts of these proteins to effect endocytosis. Overall, our results support the model that BAR domain proteins and dynamin play important roles in modulating microvesicle membrane cargo dynamics in neuroendocrine cells.

## DISCUSSION

Calcium-triggered exocytosis of SVs is a highly coordinated process involving dozens of proteins. The dynamics and assembly of these factors in live cells is not fully understood. Here, we systematically analyzed the spatial and temporal dynamics of two dozen proteins around the moment of fusion of single SLMVs in living cells. Our experiments reveal distinct local dynamics of exocytic and endocytic proteins, and a key role for BAR domain-containing proteins, along with dynamin, in modulating membrane cargo dynamics at microvesicle fusion sites.

First, we show that many proteins involved in SV exocytosis are concentrated at fusion sites several seconds before fusion (Lang *et al*., 2001; Tsuboi and Rutter, 2003; Barg *et al*., 2010; Gandasi and Barg, 2014; Ullrich *et al*., 2015; Trexler *et al*., 2016; Geerts *et al*., 2017). These include the SNAREs, VAMP2 and syntaxin1 (Figure 3), the Ca sensor synaptotagmin1 (Supplemental Figure 4), the SNARE modulators tomosyn and CAPS (Figure 4), Rab proteins, Rab3A and Rab27A, and the Rab effector molecule Rabphilin3A (Figure 2) (Supplemental Figures 2, 3 and 5). Because Rab proteins and VAMP2 are known to be associated with the vesicle membrane, their presence before fusion indicates that many SLMVs are docked at the plasma membrane prior to exocytosis. We did not measure additional recruitment of these molecules before fusion, suggesting that many proteins needed in exocytosis are pre-assembled at the vesicle. All these factors, except for synaptotagmin1, diffused away within seconds of fusion (Figures 2-4 and Supplemental Figures 2-5), indicating the highly dynamic and transient nature of the docking and fusion complex.

Synaptotagmin1 remained localized at fusion sites after fusion (Supplemental Figure 5), raising the possibility that SLMV fusion is incomplete. The complete release of VAChT-pH (Figure 1) suggests that classic kiss-and-run (Alabi and Tsien, 2013) is not the predominant mode of SLMV exocytosis in PC12 cells (Sochacki *et al*., 2012). The residual VAChT-pH signal results from VAChT trapped in neighboring clathrin structures (Sochacki *et al*., 2012). We cannot, however, rule out a cavicapture-type fusion mechanism (Holroyd *et al*., 2002; Taraska *et al*., 2003; Tsuboi *et al*., 2004), where VAChT-pH is lost, but other components such as synaptotagmin1 are retained. Previous reports have suggested that synaptotagmin stays clustered after SV exocytosis (Willig *et al*., 2006). Given that synaptotagmin1 is an important cargo for clathrin-mediated endocytosis, it is possible that it is internalized very close to fusion sites through interactions with local adaptor proteins (Haucke and De Camilli, 1999; Martina *et al*., 2001; McMahon and Boucrot, 2011). This is supported by the increase in clathrin at the plasma membrane seconds after fusion (Supplemental Figure 11).

We did not see clusters, or changes in the dynamics of SNAP25 at fusion sites (Figure 3 and Supplemental Figure 3). It is possible that high levels of endogenous SNAP25 prevent the concentration of over-expressed or labeled protein at fusion sites (Knowles *et al*., 2010). Surprisingly, we also did not see significant changes in signal for Munc18a and Munc13 (Supplemental Figures 4 and 5), molecules thought to be critical for vesicle docking and priming (Sudhof and Rothman, 2009; Gandasi and Barg, 2014), though Munc18a showed mild clustering at fusion sites (Supplemental Figure 5). This is different than the dynamics of these proteins observed with LDCVs in endocrine cells (Gandasi and Barg, 2014; Trexler *et al*., 2016). It is possible that over-expression of these proteins masked small changes in signal. But, a Munc18a construct with diminished promoter activity and reduced background signal also failed to show changes during fusion (not shown). The position and nature of the fluorescent tag could also interfere with its function (Crivat and Taraska, 2012). Detecting small transient dynamics of all proteins during fusion in this system will require future technical developments.

Of note, we found that the ATPase NSF is recruited at exocytosis (Figure 4 and Supplemental Figure 5). NSF and its binding partner α-SNAP (Sollner *et al*., 1993; Ungermann *et al*., 1998; Ryu *et al*., 2016) are thought to disassemble the SNARE complexes into monomeric SNAREs making them available for subsequent rounds of fusion (Ryu *et al*., 2016). It’s unclear if NSF acts after fusion (Littleton *et al*., 2001), or immediately prior to fusion (Banerjee *et al*., 1996; Kuner *et al*., 2008). Here, the NSF signal appears to increase at fusion sites near the beginning of fusion (Figure 4, Supplemental Figure 5), and stays elevated for ∼ 2 seconds, suggesting that NSF activity is coupled to microvesicle exocytosis, both spatially and temporally.

Importantly, we show that curvature sensing BAR domain proteins and the GTPase dynamin are locally recruited to SLMV fusion sites. Specifically, we measure dynamic recruitment of amphiphysin1, syndapin2, endophilinA1, endophilinA2 and dynamin1 to SLMVs during fusion (Figure 5 and 6, and Supplemental Figures 6, 7, and 10). Amphiphysin1 and syndapin2 recruitment was dependent on their curvature sensing BAR domains (Figure 6 and Supplemental Figure 6), suggesting that these proteins are recruited to the curved membrane of the vesicle. EndophilinB1 appeared pre-clustered at fusion sites (Supplemental Figure 7), consistent with previous findings suggesting association of endophilins with SVs at rest in nerve terminals (Bai *et al*., 2010).

Furthermore, we show that the presence of BAR domain proteins and dynamin at fusion sites slows the loss of membrane cargo following exocytosis. Supporting this idea, over-expression of endophilinA1, endophilinA2, endophilinB1, dynamin1 and dynamin2 resulted in slower VAChT decay (Supplemental Figure 7 and Figure 8). Second, amphiphysin1 mutants that failed to assemble at fusion sites resulted in faster VAChT decay (Figure 7). Third, amphiphysin1 and syndapin2 mutants that lacked the SH3 domain but showed strong recruitment to fusion sites slowed down VAChT decay (Figure 7). Fourth, dynamin1 lacking its GTPase activity, and fifth, dynamin2 deficient in binding BAR domain proteins, resulted in faster loss of VAChT at fusion sites (Figure 8). Amphiphysin1 and syndapin2 mutants lacking the SH3 domain (Figure 7) exhibited slower VAChT decay, however the prominent SH3 binding proteins dynamin1 and dynamin2 resulted in even further delays (Figure 8), and N-WASP did not show significant recruitment (Supplemental Figure 11) or affect loss of VAChT from fusion sites, suggesting that interactions with other SH3 binding proteins (McPherson *et al*., 1994; McPherson *et al*., 1996), lipids (Martin, 2015), or rearrangements in the cytoskeleton (Malacombe *et al*., 2006; Felmy, 2007; Wen *et al*., 2016; Gonzalez-Jamett *et al*., 2017) may be involved in modulating vesicle membrane cargo dynamics following fusion.

Because the loss of VAChT from fusion sites depends on both the rate of VAChT release from vesicles and the rate of VAChT diffusion in the plasma membrane, the delayed loss in the presence of BAR domain proteins and dynamins could be attributed to one or more of the following reasons. First, the recruitment of these proteins to the highly curved neck of the fusion pore could alter the structure of the pore and modulate cargo release. Such a model has been described previously for LDCVs (Wang *et al*., 2001; Llobet *et al*., 2008; Anantharam *et al*., 2010; Anantharam *et al*., 2011; Trexler *et al*., 2016). Specifically, in insulin-secreting beta cells, amphiphysin1, syndapin2, endophilinA2, dynamin1, and dynamin2 are recruited to LDCVs, and mutants of dynamin deficient in their interactions with BAR domain proteins hasten cargo release (Trexler *et al*., 2016). Moreover, in neurons, neurotransmitter release rates have been linked to the molecular machinery of the synaptic vesicle pore (Pawlu *et al*., 2004; Guzman *et al*., 2010). However, without a direct measure of microvesicle fusion pore properties in PC12 cells, it is unclear if the recruitment of BAR domain proteins modulates membrane protein release and acetylcholine release in a similar fashion. Second, BAR domain proteins and dynamin could constrict the neck of the vesicle causing endocytosis in a cavicapture-type fusion model as described for LDCVs (Holroyd *et al*., 2002; Taraska *et al*., 2003; Perrais *et al*., 2004; Taraska and Almers, 2004; Tsuboi *et al*., 2004; Fulop *et al*., 2008), or alter the merger of the omega-profile with the plasma membrane (Chiang *et al*., 2014). Examination of radial scans reveals diffusion of VAChT away from fusion sites, even in the presence of the BAR domain (Supplemental Figure 6). It is possible that most of VAChT diffuses away, and the remaining vesicle components are endocytosed, or that there is heterogeneity in the vesicle population. The third possibility is that BAR domain proteins and dynamin are not recruited to the neck of the fusion pore, but to sites very close to fusion but below the diffraction limit to effect endocytosis. This would be supported by the prolonged retention of BAR domain proteins at fusion sites (Supplemental Figure 6) and the delayed recruitment of clathrin (Supplemental Figure 11), and is consistent with previous studies demonstrating spatial and temporal proximity of exocytosis and compensatory endocytosis (Roos and Kelly, 1999; Sochacki *et al*., 2012; Watanabe *et al*., 2013a; Watanabe *et al*., 2013c; Wu *et al*., 2014). And lastly, BAR domain proteins and dynamin may have non-local effects on plasma membrane tension (Stachowiak *et al*., 2012), the cytoskeleton, or membrane fluidity, and affect fusion dynamics and/or diffusion of membrane cargo without directly modulating the fusion pore or endocytosis.

At the neuronal synapse, the association of endocytic proteins with SVs is thought to ensure the availability of these proteins for compensatory endocytosis (Shupliakov, 2009; Bai *et al*., 2010). Our findings that endocytic proteins are dynamically recruited to sites of fusing SLMVs, which are structurally and functionally similar to SVs (Thomas-Reetz and De Camilli, 1994), suggests that these proteins could play more active roles in modulating exocytosis, endocytosis, and the coupling between these two processes. Direct evaluation of the effects of endocytic protein recruitment on fusion pore dynamics and rapid compensatory endocytosis in neurons could provide insights into the complex regulation of synaptic membrane trafficking.

The protein dynamics observed at SLMV fusion sites is like that seen with the much larger LDCVs in endocrine cells. While SVs and LDCVs have broadly similar molecular requirements for fusion (Xu and Xu, 2008), differences in their size (∼ 50 nm vs. ∼ 150 nm diameter), Ca^2+^ dependencies (Verhage *et al*., 1991; Heidelberger *et al*., 1994; Ninomiya *et al*., 1997; Sudhof, 2013a), and latencies to fusion (Chow *et al*., 1992; Sabatini and Regehr, 1996), suggest differential regulation of the exocytic fusion machinery. Furthermore, their different curvatures, pore sizes (19 pS vs 213 pS conductance) (Klyachko and Jackson, 2002), and pore stabilities (Zhang and Jackson, 2010) suggest distinct structures of the pore. However, in both SLMVs and LDCVs, syntaxin1 (Lang *et al*., 2001; Barg *et al*., 2010; Gandasi and Barg, 2014), VAMP (Tsuboi and Rutter, 2003; Trexler *et al*., 2016), Rab3A (Gandasi and Barg, 2014; Trexler *et al*., 2016), Rab27A, Rabphilin3A, CAPS and tomosyn (Trexler *et al*., 2016) are concentrated at fusion sites, and diffuse away following fusion, supporting parallels in the molecular assembly and disassembly of key components during LDCV and microvesicle exocytosis. Moreover, BAR domain proteins and dynamin are recruited to fusion sites of both SLMVs and LDCVs sites (Holroyd *et al*., 2002; Tsuboi *et al*., 2004; Trexler *et al*., 2016). Given the different sensitivities to membrane curvature among the various BAR domain proteins (Daumke *et al*., 2014) and dynamin isoforms (Yoshida *et al*., 2004; Liu *et al*., 2011), the differences in their assembly and effects on LDCV and SLMV exocytosis may be nuanced. For example, all three endophilins associated with fusing SLMVs, but only endophilinA2 localizes with LDCVs (Trexler *et al*., 2016). Future work at higher temporal or spatial resolutions comparing LDCV and microvesicle fusion in the same cells, under expression-controlled conditions, will help identify key differences in protein dynamics needed to regulate different pools of vesicles, their fusion kinetics, release of their unique cargos, and their roles in signaling and cellular communication.

## MATERIALS AND METHODS

### Cells and solutions

PC12-GR5 cells were grown in DMEM containing 4 mM L-glutamine, supplemented with 5 % fetal bovine serum, 5 % horse serum, and 1 % penicillin/streptomycin, at 37 °C in 5% CO_2_. The cell-line was originally obtained from Rae Nishi (MBL), expanded from low passage frozen stocks, and was not further authenticated. The cells tested negative for mycoplasma contamination. Cells were plated onto 25-mm, #1.5, round, poly-D-lysine coated glass cover-slips, and transfected approximately 24–48 h later with 1 µg each of the indicated plasmids, or 25 nmol of siRNA (Dharmacon) using Lipofectamine 2000 (Invitrogen) following the manufacturer’s protocol. The list of plasmids used and the n-values for all the experiments are in Table 1. Cells were imaged approximately 24-48 h after transfection. The imaging buffer contained (in mM): 130 NaCl, 2.8 KCl, 5 CaCl_2_, 1 MgCl_2_, 10 HEPES and 10 glucose. pH was adjusted to 7.4 with 1 N NaOH. The stimulation buffer contained (in mM): 50 NaCl, 105 KCl, 5 CaCl_2_, 1 MgCl_2_, 10 HEPES and 1 NaH_2_PO_4_. pH was adjusted to 7.4 with 5 M KOH.

### TIRF microscopy

TIRF microscopy was done as previously described (Trexler et al, 2016; Trexler and Taraska, 2017). Cells were imaged on an inverted fluorescent microscope (IX-81, Olympus), equipped with a X100, 1.45 NA objective (Olympus). Combined green (488 nm) and red (561 nm) lasers (Melles Griot) were controlled with an acousto-optic tunable filter (Andor), and passed through a LF405/488/561/635 dichroic mirror. Emitted light was filtered using a 565 DCXR dichroic mirror on the image splitter (Photometrics), passed through 525Q/50 and 605Q/55 filters, and projected onto the chip of an EM-CCD camera. Images were acquired using the Andor IQ2 software. Cells were excited with alternate green and red excitation light, and images in each channel were acquired at 100 ms exposure, at 5 Hz. To trigger exocytosis, stimulation buffer was applied for 30 – 40 s using a µFlow perfusion system (ALA Scientific Instruments) with a 100 µm pipette positioned close to the surface of the cell. Each day before experiments, 100 nm yellow-green fluorescent beads (Invitrogen) were imaged in the green and red channels, and superimposed by mapping corresponding bead positions. The green and red cell images were aligned post-acquisition using projective image transformation as described before (Sochacki et al, 2012; Trexler et al, 2016). All experiments were carried out at 25 °C.

### Image analysis

Image analysis was performed using Metamorph (Molecular Devices), ImageJ (NIH) and custom scripts on MATLAB (Mathworks). The co-ordinates of the brightest pixel in the first frame of brightening for individual fusion events in the green channel were identified by eye, and assigned as the center of the fusion event. All events were time-aligned to the first frame of fusion (0 seconds). A circular ROI of 6 pixels (∼ 990 nm) diameter and a square of 21 pixels (∼3.5 µm) were drawn around the center, and the mean intensities in the surrounding square, excluding the circular region, were subtracted from the mean intensities in the center for each fusion event. This analysis was done for every frame from – 10 s to + 50 s relative to the first frame of fusion. The background subtracted time-lapse intensities in the green channel were normalized 0 to 1 for each event, with ‘0’ being the mean pre-fusion value, obtained from −5 s to −3 s, and ‘1’ being the peak intensity. The background subtracted time-lapse intensities for each event in the red channel were also normalized 0 to 1, with ‘0’ being the minimum, and ‘1’ the maximum value, within – 10 s to + 50 s. The resulting normalized traces were averaged across all events, and truncated to show data from −5 seconds to +10 seconds to better represent the protein dynamics around the moment of fusion. For the radial scan analyses, the intensities along 32 overlapping lines of 3.5 µm length arranged in a star pattern were averaged for each frame. The resulting kymograph through time for each event was normalized from 0 to 1000, and averaged across all events. Local background subtraction was not performed for radial scan analyses. The VAChT decay constants were obtained by single exponential fits of average VAChT fluorescence from 0.2 s to 5 seconds in the time-lapse traces, with y-offset constrained to 0, using OriginPro. Fusion events were excluded from analysis if one or more additional fusion events occurred in the circular ROI within 5 seconds before or after fusion (−5 s to 5 s in the time-lapse traces). If additional events occurred > 5 s after fusion, the original fusion event was not excluded from analysis. Therefore, the average intensities of VAChT and red proteins after 5 s in traces and radial scan images should be interpreted with this caveat in mind.

### Western Blots

PC12 cells were transfected with Syndapin2-GFP, or siRNA (Dharmacon) using Lipofectamine 2000, and protein was isolated ∼ 24 h later. Cells were lysed in RIPA buffer (1 % Anatrace, 0.5 % sodium deoxycholate, 0.1 % SDS, 2mM EDTA and 1mM phenylmethylsulfonyl fluoride in PBS) on ice for 30 min. Lysates were centrifuged at 12,000g at 4°C for 10 min. Supernatants were boiled in lithium dodecyl sulphate (LDS) sample buffer containing 62.5 mM dithiothreitol for 10 min, and loaded onto 4-12 % Tris-Bis gels (NuPAGE). Protein was transferred onto nitrocellulose membrane using the iBlot dry transfer system (ThermoFisher Scientific), and syndapin2 detected with monoclonal antibody (ThermoFisher Scientific), and peroxidase-labelled secondary antibody. Blots were stripped with stripping buffer (ThermoFisher Scientific), and re-probed with α-actin antibody (Abcam).

### Statistical Tests

All data are expressed as mean ± SEM. Statistical analysis was done using two-tailed paired Student’s t-test to compare normalized red protein fluorescence intensities before and after fusion for the same set of events, as indicated in figure legends. *p* < 0.01 was used as a measure of statistical significance.

## ACKNOWLEDGEMENTS

We thank Marie-Paule Strub (NIH) for assistance with molecular biology, J. Silver, K. Sochacki and A. Trexler for critical reading of the manuscript, and J. Silver, S. Kale and members of the Taraska laboratory for discussions and input on data analysis methods. J.W.T. is supported by the Intramural Research Program of the National Heart, Lung, and Blood Institute, National Institutes of Health.

## AUTHOR CONTRIBUTIONS

A.S. and J.W.T. designed study, A.S. performed experiments, A.S. and J.W.T. analyzed data and wrote the manuscript.

---

**Supplemental Figure 1.**
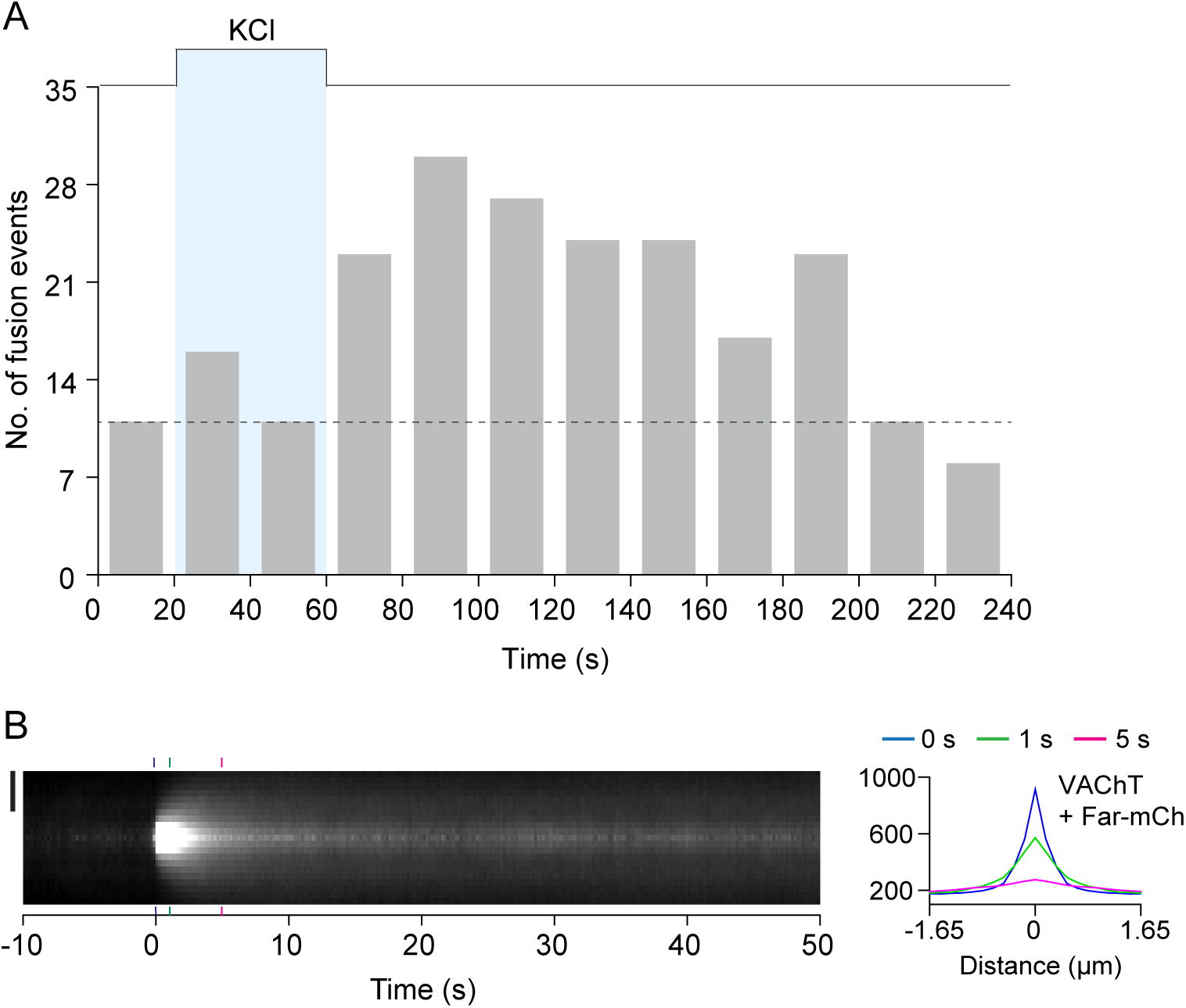
Frequency of microvesicle fusion events, and radial spread of VAChT-pH following exocytosis, in PC12 cells. (A) Histogram showing number of fusion events before and after application of stimulation buffer (KCl). (B) Average normalized radial line scans of VAChT-pH during fusion. Contrast has been adjusted to show areas with lower intensity. Vertical dotted lines indicate the time-points corresponding to linescan intensity plots (Right). Standard errors are plotted as shaded areas around the average traces. Data was obtained from cells co-expressing VAChT-pH and farnesyl-mCherry.

**Supplemental Figure 2.**
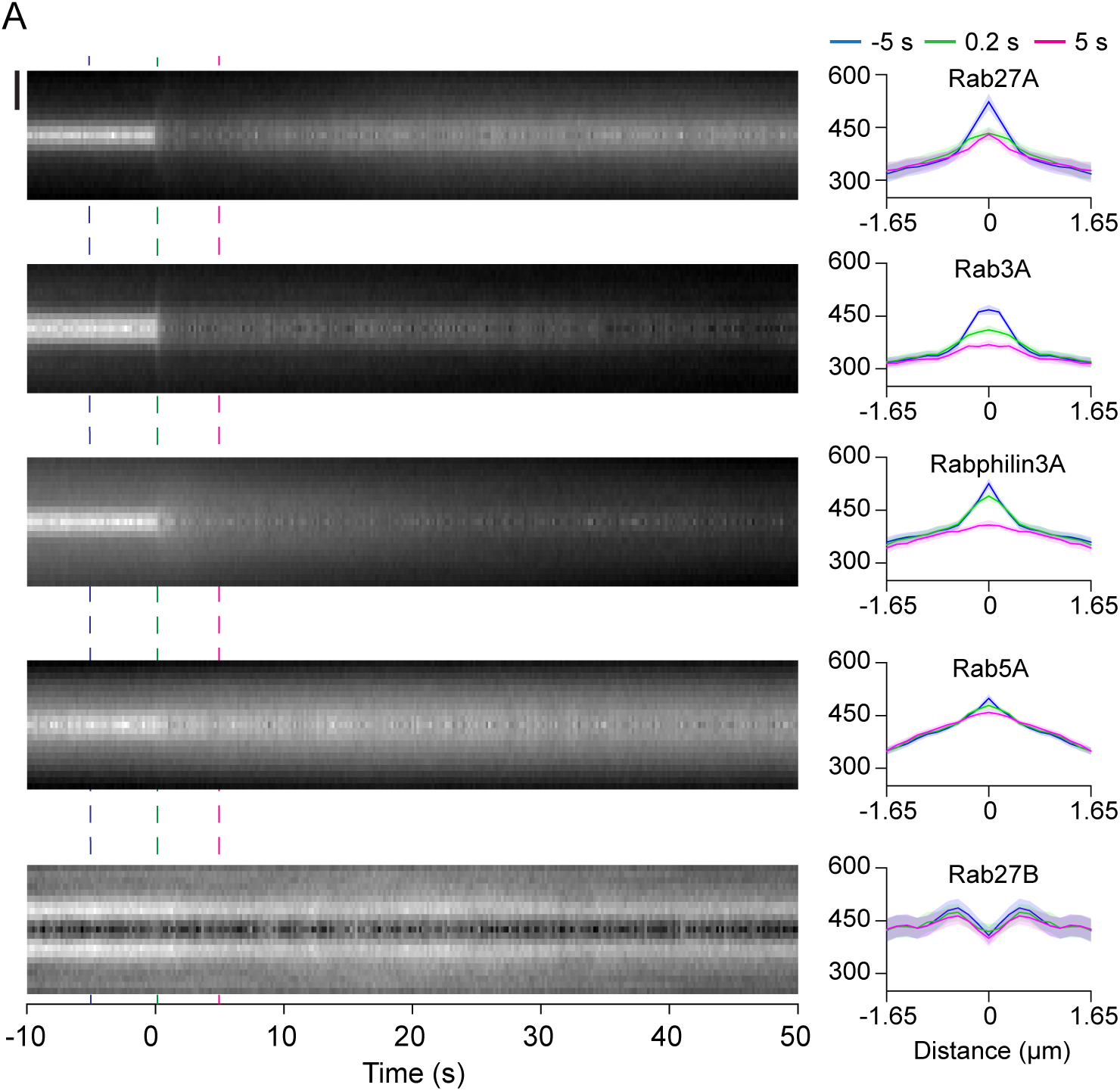
Dynamics of Rab proteins during SLMV fusion. (A) Average normalized radial line scans of the Rab proteins during microvesicle exocytosis. Bar, 1 μm. Vertical dotted lines indicate the time-points corresponding to linescan intensity plots (Right). Standard errors are plotted as shaded areas around the average traces.

**Supplemental Figure 3.**
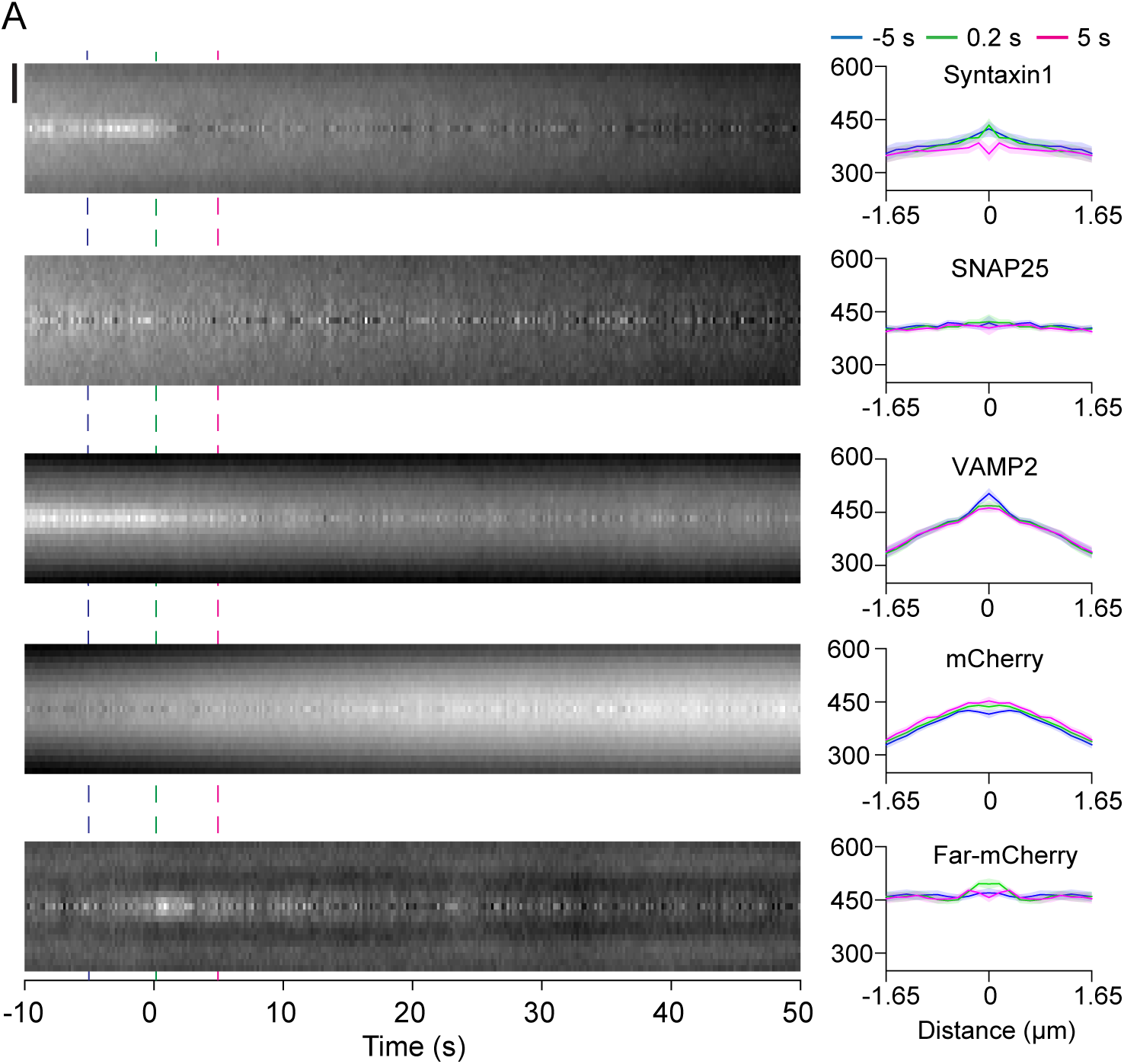
Dynamics of SNARE proteins during fusion. (A) Average normalized radial line scan analysis of SNAREs and control mCherry proteins during microvesicle exocytosis. Bar, 1 μm. Vertical dotted lines indicate the time-points corresponding to linescan intensity plots (Right). Standard errors are plotted as shaded areas around the average traces.

**Supplemental Figure 4.**
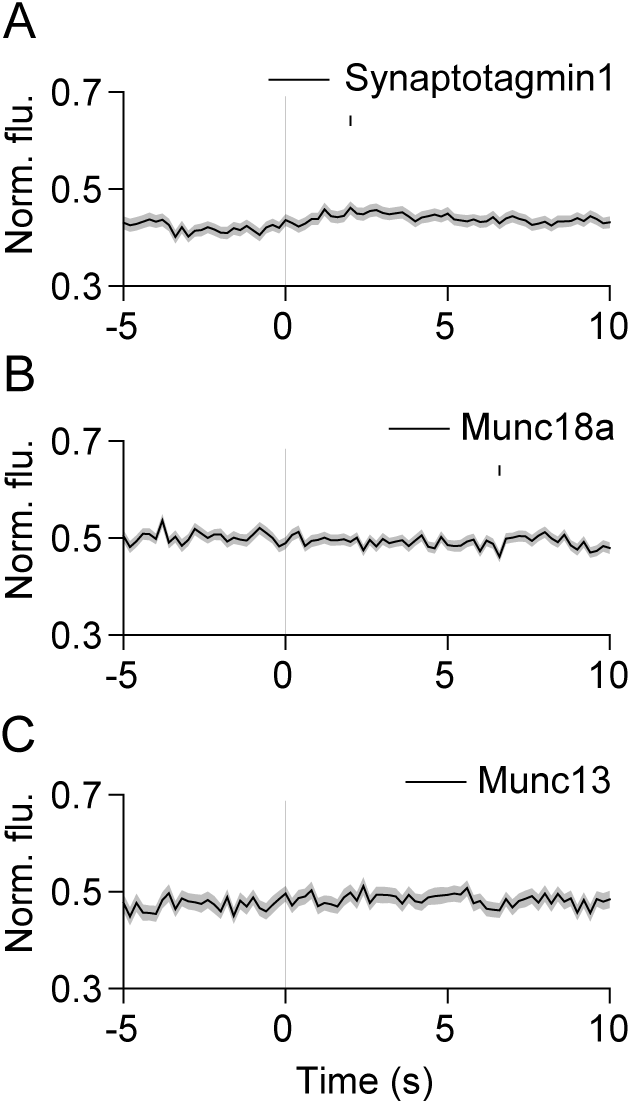
Synaptotagmin1 and Munc protein behavior at SLMV fusion sites. (A-C) Average time-lapse traces of normalized fluorescence intensities for: (A) synaptotagmin1-mCherry (266 events, 5 cells), (B) mCherry-Munc18a (250 events, 11 cells), and (C) Munc13-mCherry (132 events, 5 cells). Individual event traces were time-aligned to 0 s (vertical black line), which corresponds to the fusion frame in the green channel. Small vertical black lines indicate p < 0.05 (paired Student’s t-test) when comparing every time-point after 3 s with the average pre-fusion value obtained from −5 to −3 s. Standard errors are plotted as shaded areas around the average traces.

**Supplemental Figure 5.**
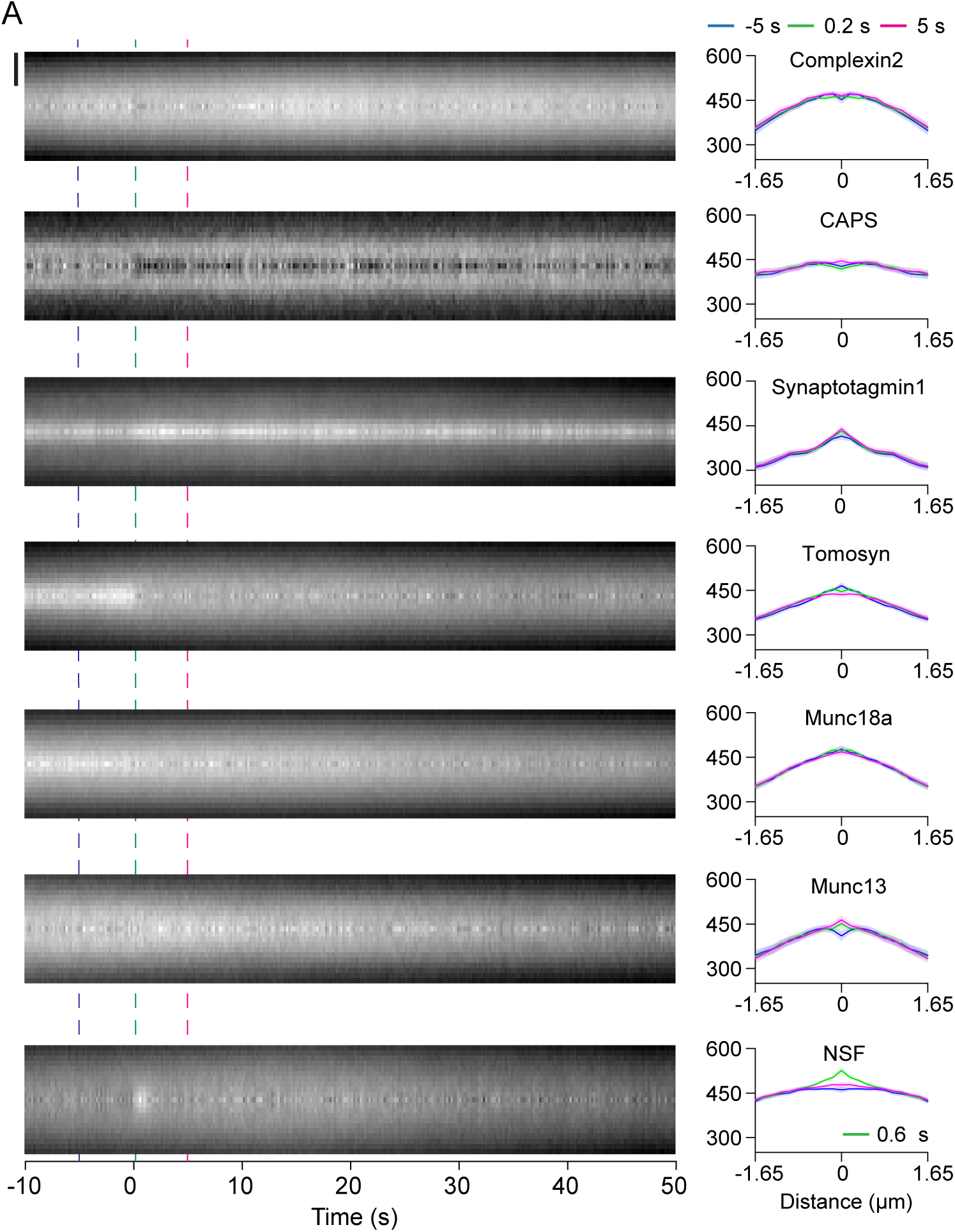
Dynamics of SNARE modulators during SLMV fusion. (A) Average normalized radial line scans of accessory proteins during microvesicle exocytosis. Bar, 1 μm. Vertical dotted lines indicate the time-points corresponding to linescan intensity plots (Right). Standard errors are plotted as shaded areas around the average traces.

**Supplemental Figure 6.**
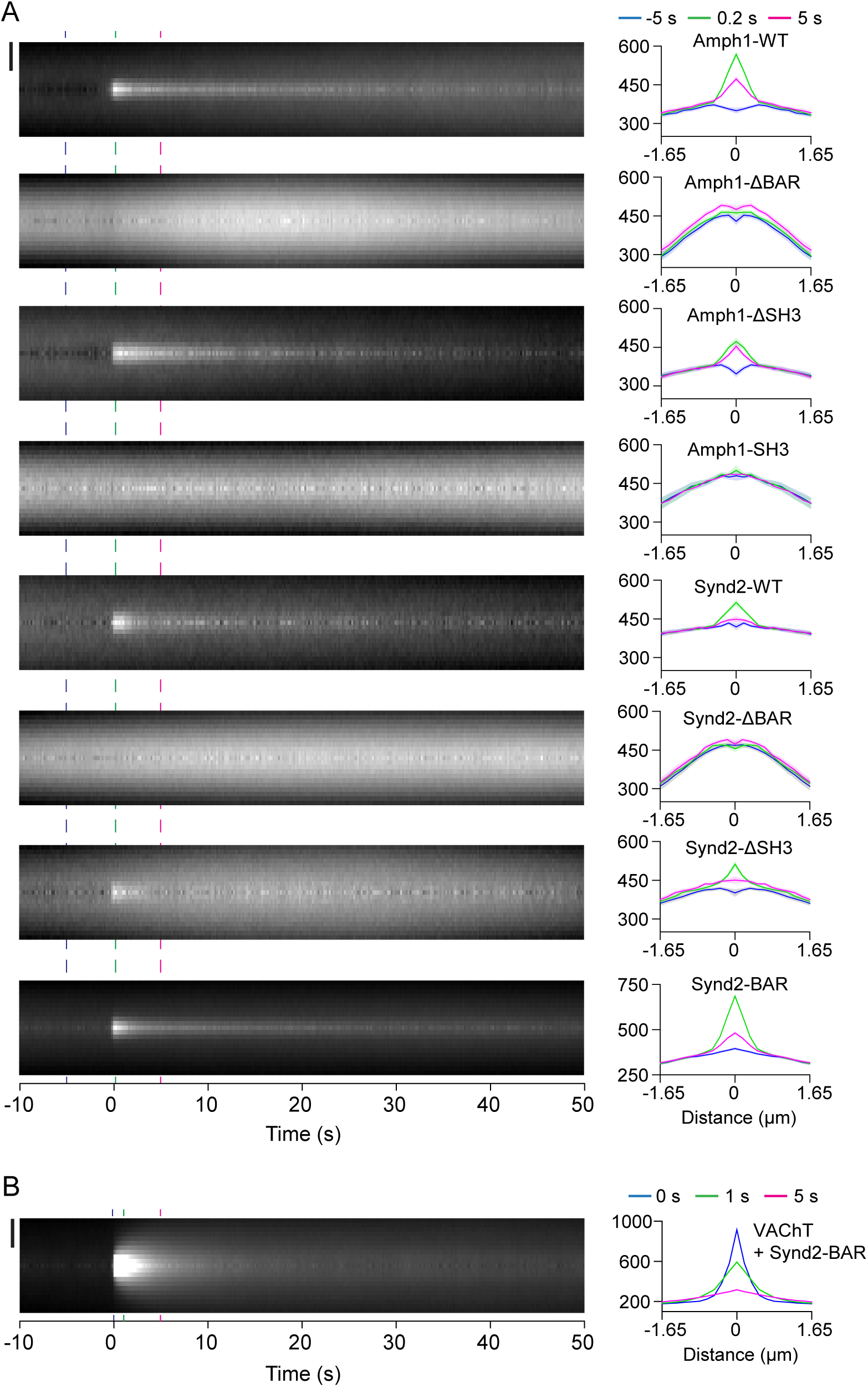
Dynamics of WT and mutant amphiphysin1 and syndapin2, and diffusion of VAChT-pH, during SLMV fusion. (A) Average normalized radial line scans of WT and mutant amphiphysin1 and syndapin2 proteins during microvesicle exocytosis. Bar, 1 μm. (B) Average normalized radial line scans of VAChT-pH during fusion in PC12 cells co-expressing syndapin2-BAR. Contrast has been adjusted to show areas with lower intensity. Vertical dotted lines indicate the time-points corresponding to linescan intensity plots (Right). Standard errors are plotted as shaded areas around the average traces.

**Supplemental Figure 7.**
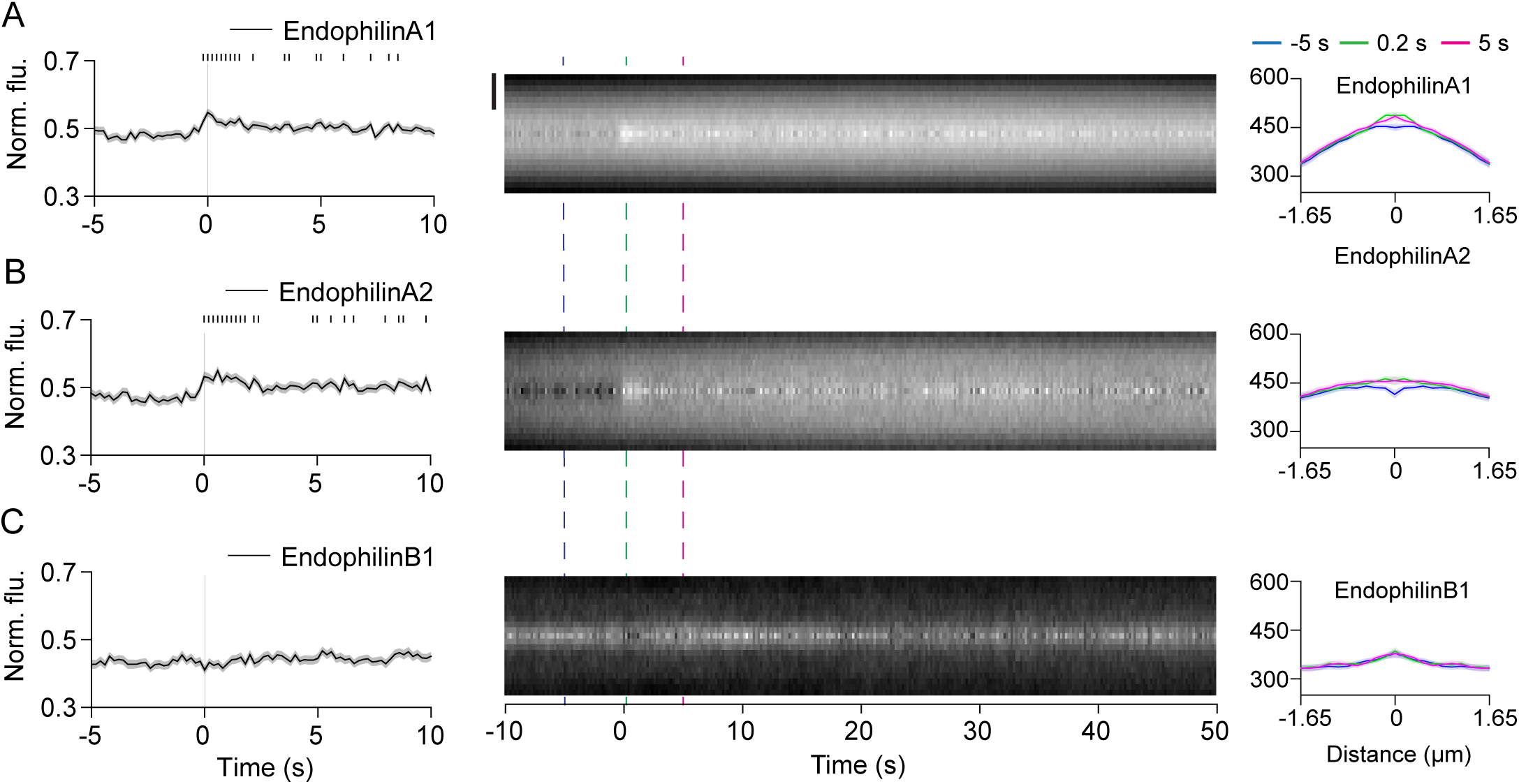
Endophilins are localized at SLMV fusion sites during exocytosis. (A-C) (Left) Average time-lapse traces of normalized fluorescence intensities for: (A) endophilinA1-mCherry (258 events, 13 cells), (B) endophilinA2-mCherry (207 events, 11 cells), and (C) endophilinB1-mCherry (213 events, 11 cells). Individual event traces were time-aligned to 0 s (vertical black line), which corresponds to the fusion frame in the green channel. Small vertical black lines indicate p < 0.05 (paired Student’s t-test) when comparing every time-point after 3 s with the average pre-fusion value obtained from −5 to −3 s. (Middle) Average normalized radial line scans of endophilins during fusion. Bar, 1 μm. Vertical dotted lines indicate the time-points corresponding to linescan intensity plots (Right). Standard errors are plotted as shaded areas around the average traces.

**Supplemental Figure 8.**
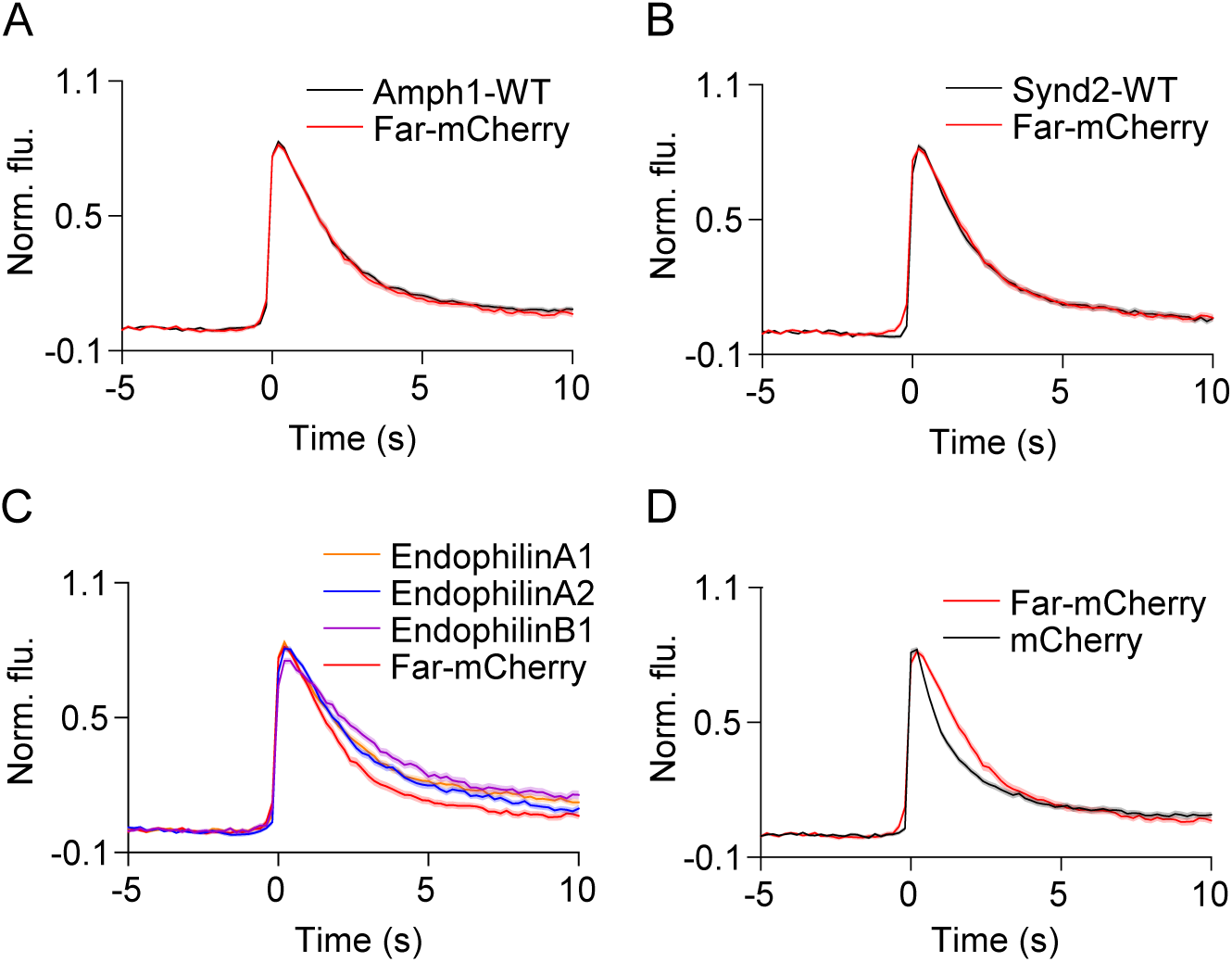
Effect of BAR domain proteins on loss of VAChT-pH from SLMV fusion sites. (A-D) Average time-lapse traces of normalized VAChT-pH fluorescence intensities in cells co-expressing VAChT-pH and the indicated BAR domain or control mCherry proteins. Individual traces were time-aligned to 0 s, which corresponds to the frame of fusion. Standard errors are plotted as shaded areas around the average traces.

**Supplemental Figure 9.**
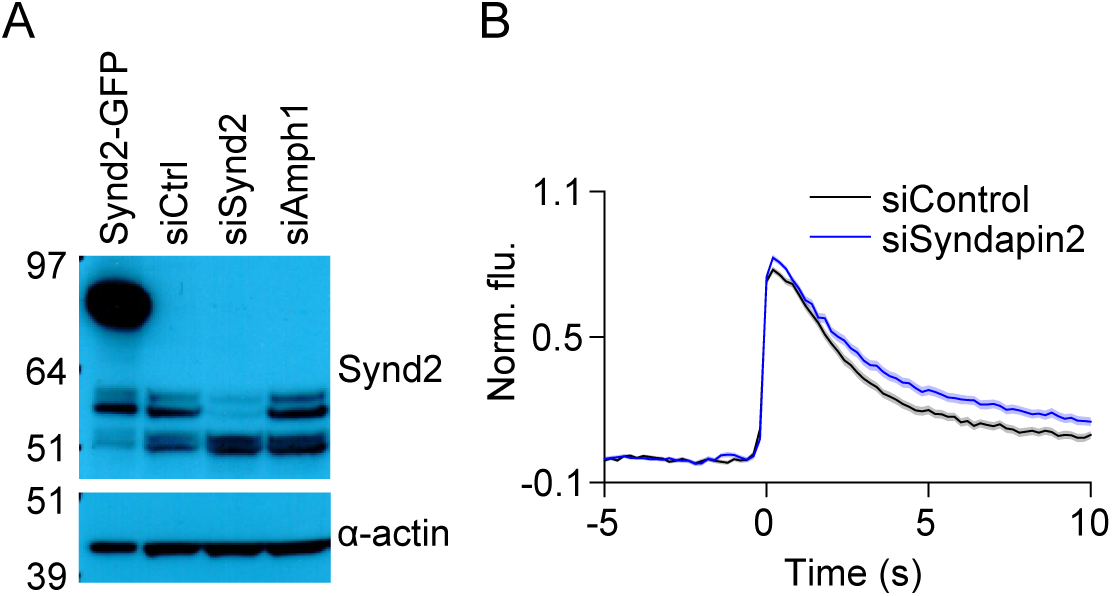
Effect of syndapin2 knock-down on VAChT-pH loss from fusion sites. (A) Western blots showing expression of syndapin2 in PC12 cells, that is reduced following treatment with siRNA. (Top) Blot probed with anti-syndapin2 antibody. (Bottom) Same blot re-probed with anti-α-actin. (B) Average time-lapse traces of normalized VAChT-pH fluorescence intensities in cells treated with control siRNA (219 events, 15 cells) or sisyndapin2 (184 events, 16 cells). Individual traces were time-aligned to 0 s, which corresponds to the fusion frame. Standard errors are plotted as shaded areas around the average traces.

**Supplemental Figure 10.**
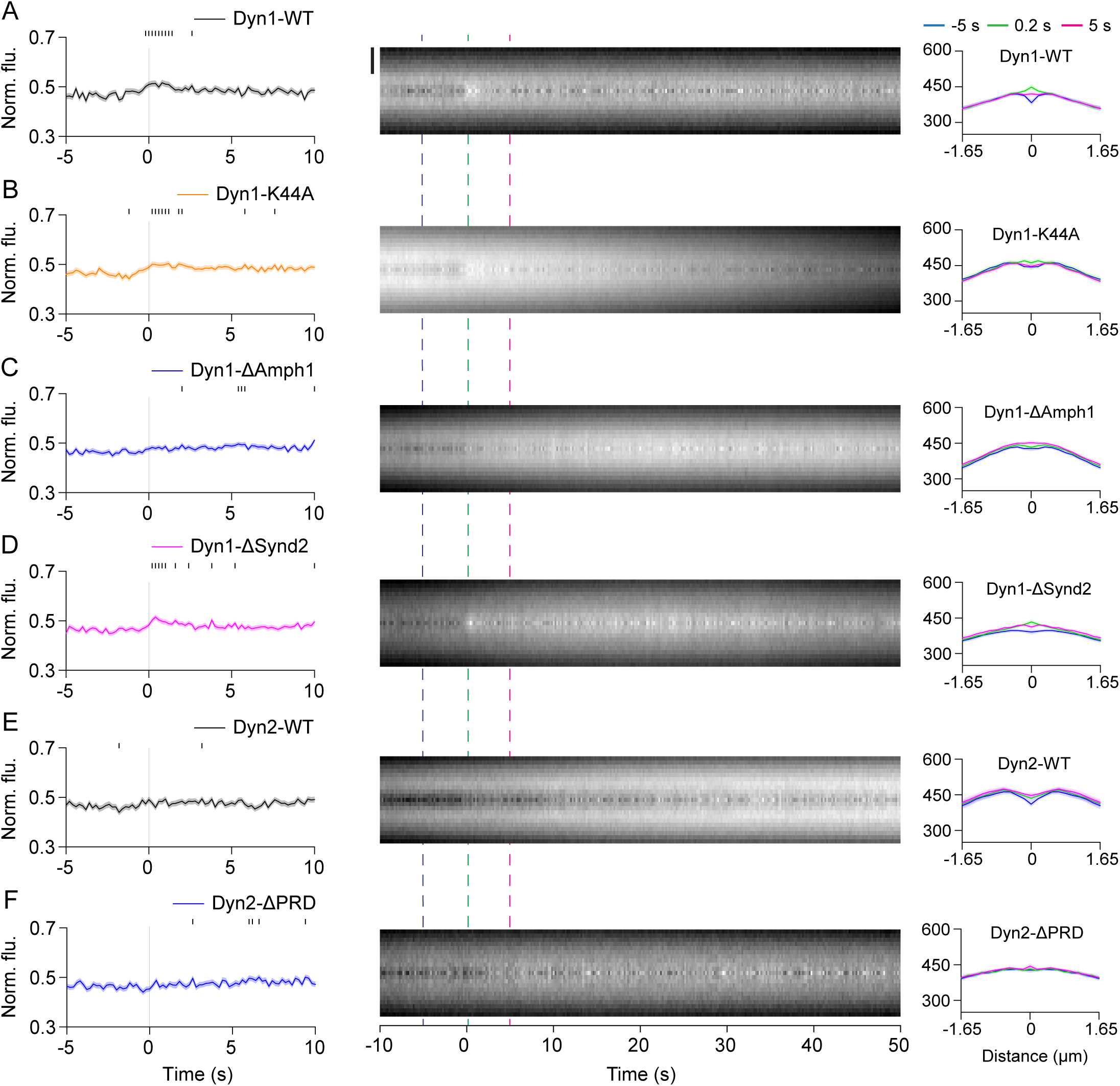
Recruitment of WT and mutant dynamin to SLMV fusion sites. (A-F) (Left) Average time-lapse traces of normalized fluorescence intensities for: (A) Dynamin1-WT (250 events, 14 cells), (B) Dynamin-K44A (309 events, 12 cells), (C) Dynamin1-ΔAmph1 (365 events, 17 cells), (D) Dynamin1-ΔSynd2 (341 events, 17 cells), (E) Dynamin2-WT (260 events, 12 cells), and (F) Dynamin2-ΔPRD (252 events, 14 cells). Individual event traces were time-aligned to 0 s (vertical black line), which corresponds to the fusion frame in the green channel. Small vertical black lines indicate p < 0.05 (paired Student’s t-test) when comparing every time-point after 3 s with the average pre-fusion value obtained from −5 to −3 s. (Middle) Average normalized radial line scans of WT and mutant dynamin proteins during microvesicle exocytosis. Bar, 1 μm. Vertical dotted lines indicate the time-points corresponding to linescan intensity plots (Right). Standard errors are plotted as shaded areas around the average traces.

**Supplemental Figure 11.**
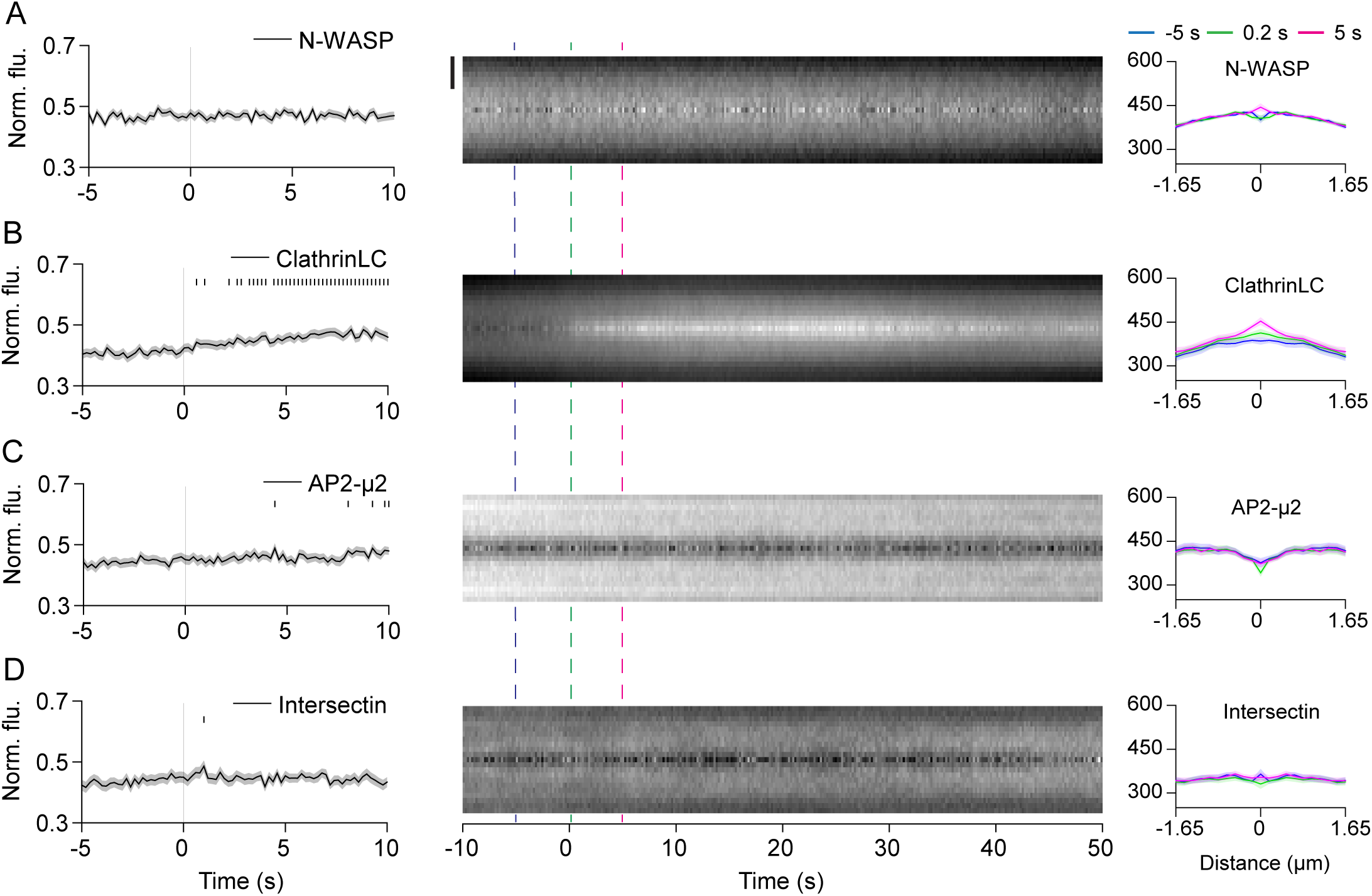
Dynamics of other endocytic proteins at SLMV fusion sites. (A-D) (Left) Average time-lapse traces of normalized fluorescence intensities for: (A) N-WASP-mCherry (197 events, 8 cells), (B) ClathrinLC-mCherry (209 events, 5 cells), (C) AP2-µ2-mCherry (153 events, 5 cells), and (D) mCherry-Intersectin (122 events, 5 cells). Individual event traces were time-aligned to 0 s (vertical black line), which corresponds to the fusion frame in the green channel. (Middle) Average normalized radial line scan analysis. Bar, 1 μm. Vertical dotted lines indicate the time-points corresponding to linescan intensity plots (Right). Standard errors are plotted as shaded areas around the average traces.

**Supplemental Figure 12.**
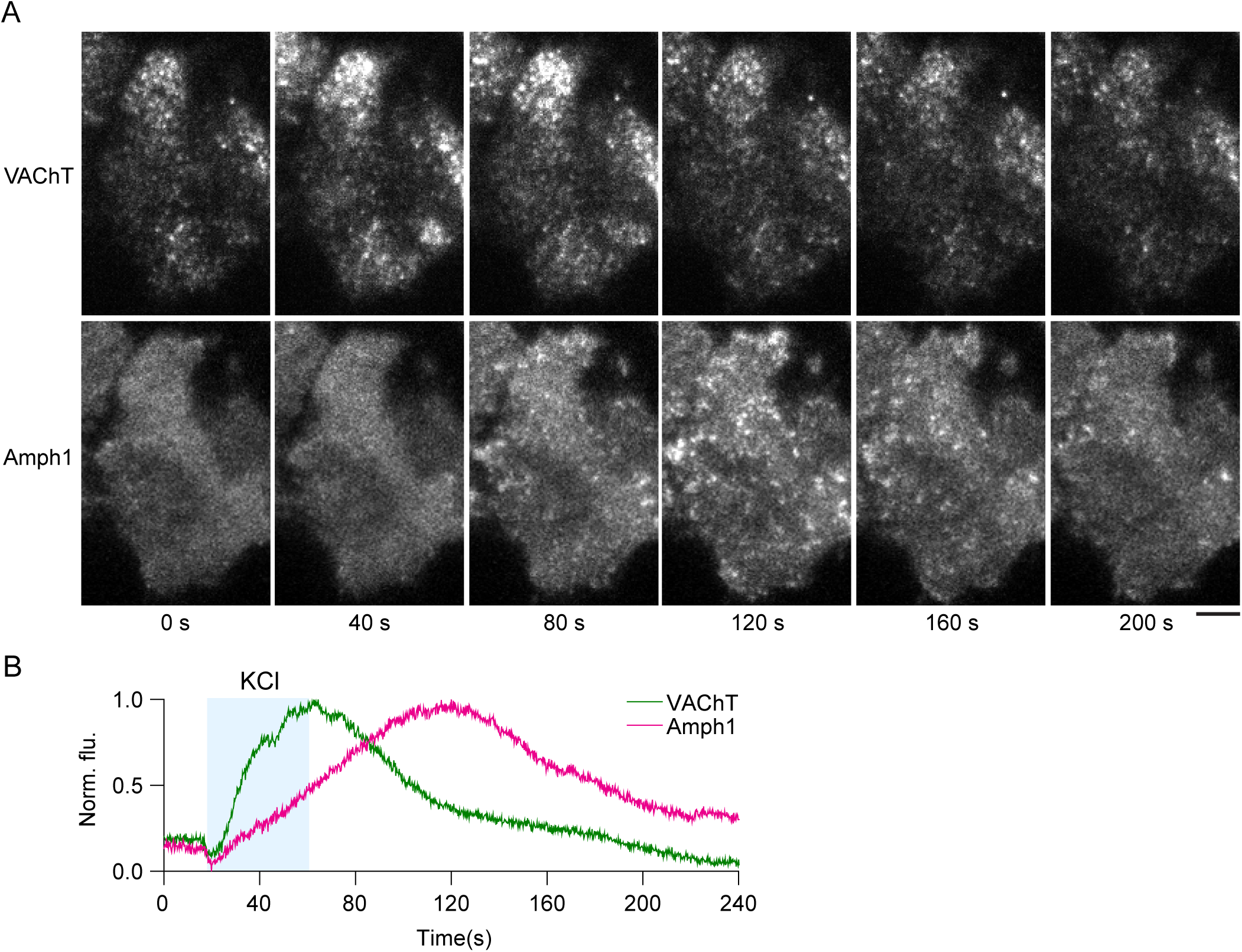
Plasma membrane recruitment of amphiphysin1 following stimulation. (A) Images of a PC12 cell expressing VAChT-pH (top) and Amph1-mCherry (bottom) at indicated time-points, with ‘0 s’ being the start of the experiment. Stimulation buffer (KCl) was applied from ∼ 20 s to 60 s. Bar, 5 μm. (B) Time-lapse traces show background-subtracted and normalized mean fluorescence intensities for VAChT-pH (green) and amphiphysin1-mCherry (magenta) from a region around the whole cell.

